# *Breakthrough*: A First-In-Class Virtual Simulator for Dose Optimization of ACE Inhibitors in Veterinary Cardiology

**DOI:** 10.1101/2022.11.14.516497

**Authors:** Benjamin K. Schneider, Jessica Ward, Samantha Sotillo, Catherine Garelli-Paar, Emilie Guillot, Marc Prikazsky, Jonathan P. Mochel

## Abstract

The human and canine renin-angiotensin-aldosterone-systems (RAAS) play a central role in the pathophysiology of congestive heart failure (CHF), justifying the use of angiotensin converting enzyme inhibitors inhibitors (ACEi) in this indication. Seminal studies in canine CHF had suggested that the pharmacological action of benazepril was relatively independent of doses > 0.25 mg/kg P.O, thereby providing a rationale for the European label dose of 0.25 mg/kg P.O q24h in dogs with cardiovascular diseases. However, most of these earlier studies on benazepril pharmacodynamics relied on measures of ACE activity – a sub-optimal endpoint to characterize the effect of benazepril on the RAAS.

Nonlinear mixed-effects (NLME) modeling is an established framework for characterizing the effect of therapeutics on complex biological systems, such as the RAAS cascade. Importantly for therapeutic schedule optimization, one can use such a model to predict the outcomes of various hypothetical dosing schedules via simulation.

The objectives of this study were <u>**(i)**</u> to expand on previous NLME modeling efforts of the dose-exposure-response relationship of benazepril on biomarkers of the RAAS which are relevant to CHF pathophysiology and disease prognosis {angiotensins I, II, III, IV, (1-7)} by using a quantitative systems pharmacology (QSP) modeling approach; and <u>**(ii)**</u> to develop a software implementation of the model capable of simulating clinical trials in benazepril in dogs bedside dose optimization.

This study expands on previous modeling efforts to characterize the changes in RAAS pharmacodynamics in response to benazepril administration and showcase how QSP modeling can be used for efficient dose optimization of ACEis at the bedside. Our results suggest that 0.5 mg/kg PO q12h of benazepril produced the most robust reduction in AngII and upregulation of RAAS *alternative pathway* biomarkers. This model will eventually be expanded to include relevant clinical endpoints, which will be evaluated in an upcoming prospective trial in canine patients with CHF.

**Author Summary:** Congestive heart failure (CHF) is a disease of the heart, common to both dogs and humans, where the heart is not healthy enough to pump blood around the body efficiently. Because the blood isn’t moving around the body as efficiently, it tends to get congested in various areas of the body and increases strain on the heart. Benazepril is a drug for CHF used in both dogs and humans to reduce congestion and improve the functioning of the cardiovascular system. Although benazepril is effective, there’s evidence that suggests the dosing could be improved if the therapeutic was further studied.

In this experiment, we tested benazepril at several safe dosages in well-cared for and healthy dogs to collect data on the relationship between dose size, dosing frequency, and effect on the cardiovascular system. Using this data, we built computer models of benazepril to simulate many clinical trials. By studying these simulations, we were able to make several predictions about the optimal dosing schedule of benazepril in dogs. We’ve also built a web-app version of the computer model for veterinary researchers to use, modify, and study. This work also provides a platform and roadmap for optimizing benazepril dosages in human CHF.

## Introduction

Although the exact pathophysiology of the heart diseases underlying congestive heart failure (CHF) differ between man and his best friend, overactivation of the renin-angiotensin-aldosterone system (RAAS) plays a key role in the pathogenesis and development of CHF in both humans and dogs. To reduce RAAS activation, there is a substantial history of using ACEis, such as benazepril, to treat CHF in both species (1–3). This makes the use of benazepril to treat CHF in canines and humans an excellent case study for applying the *One Health Initiative* paradigm. This paradigm recognizes that accumulating data on the effect of therapeutics on CHF in canines has the potential to benefit therapeutic management of CHF in humans and *vice versa* (4).

The RAAS is a neurohormonal compensatory system which primarily manages blood volume and pressure by modulating electrolyte transport and vascular tone. The contemporary model of RAAS activation has two main components. The *classical* RAAS pathway refers to the peptide cascade from angiotensinogen to angiotensin I (AngI), and then from AngI to angiotensin II (AngII). These enzymatic reactions are catalyzed by renin and ACE, respectively, and ultimately lead to increased aldosterone (ALD) production (see **Fig. 1**; (4)). Short-term physiologic consequences of *classical* RAAS activation include vasoconstriction, renal sodium and water retention, and increased blood pressure. Long-term physiological consequences include fluid overload, increased cardiac afterload, and myocardial and vascular fibrosis (5–9). Essentially, chronic long-term *classical* RAAS activation both contributes to, and is stimulated by, the development of CHF (10,11), while *classical* RAAS pathway downregulation has been associated with improved long-term prognosis in CHF (9,12–15). The *alternative* RAAS pathway acts as a counterregulatory mechanism against *classical* pathway activation. Activation of the *alternative* RAAS pathway is characterized by catalysis of AngII to angiotensin (1-7) (i.e. Ang(1-7)) by the enzyme ACE2. In turn, Ang(1-7) activates Mas receptors leading to vasodilatation, diuresis, and natriuresis (16). Via this physiological effect, chronic *alternative* RAAS activation in CHF has been associated with reduced risk of heart failure in patients with reduced and preserved ejection fraction. An ideal therapeutic drug candidate for CHF would therefore modulate both pathways at once, downregulating the activity of the *classical* RAAS while preserving or upregulating the *alternative* RAAS pathway (13). However, little is known about the effect of benazepril on the *alternative* RAAS in either humans or dogs.

**Figure 1:**
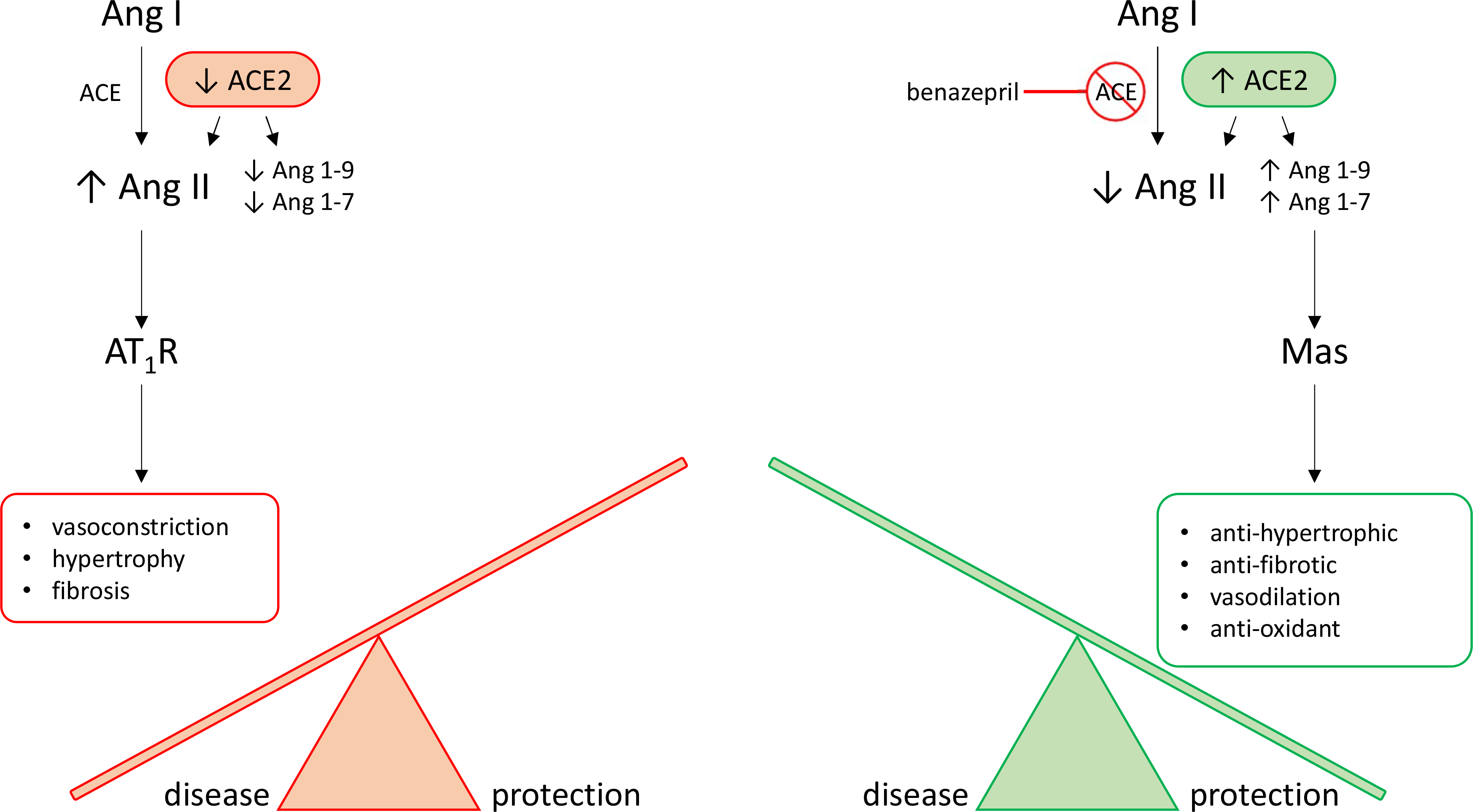
Arms of the RAAS. RAAS activation is thought of as having two main pathways which act as counterregulatory mechanisms for one another. The <u>classical RAAS</u> pathway (in red-orange) refers to the peptide cascade from angiotensin I (Ang I) to angiotensin II (Ang II) via ACE. This stimulates aldosterone production which then activates AT_1_ receptors (AT_1_R). Physiologic consequences of classical RAAS activation, including vasoconstriction, hypertrophy, and fibrosis, typical worsen congestive heart failure (CHF). Benazepril inhibits ACE, therefore activating the <u>alternative RAAS</u> pathway (in green). Activation of the alternative RAAS pathway is characterized by catalysis of Ang II to Ang1-7 by the enzyme ACE2. In turn, Ang1-7 activates Mas receptors leading to vasodilatation, diuresis, and natriuresis. These effects are protective against CHF. Our goal is to use mathematical modeling to determine a dosage which both reduces classical RAAS pathway activation and stimulates alternative RAAS pathway activation. This hypothetical dosage would maximize CHF-protective effects of benazepril.

Benazepril hydrochloride is a non-sulfhydryl ACEi commonly used for the management of CHF in both humans and dogs. Like other ACEi, benazepril is a prodrug that is rapidly converted through hydrolysis to its active benazeprilat by esterases, mainly in the liver (17). Although frequently prescribed, the recommended dosage range of benazepril is quite broad and there is no clear consensus on the ideal dose to be used in patients with CHF. In humans, benazepril is typically prescribed for hypertension at an initial dose of 2.5-10 mg per day and up titrated to 20 or 40 mg per day, administered either once or twice daily (q24h or q12h) which is roughly equivalent to 0.5 mg/kg q12h for a 60 kg adult (18). In dogs, the labeled dose of benazepril in the EU is 0.25-1.0 mg/kg PO q24h, whereas ACVIM veterinary consensus statements recommend a dose of 0.5 mg/kg PO q12h (19). Pharmacokinetic (PK) and pharmacodynamic (PD) studies comparing various doses of benazepril in healthy dogs have not provided consistent recommendations to date. The study that was used for registration of benazepril in the EU showed that a single PO dose of benazepril effectively suppressed ACE activity for up to 24 hours, and that ACE inhibition in plasma was independent of dosage ≥ 0.25 mg/kg (2). However, subsequent reanalysis of these data using mathematical modeling suggested that q12h dosing (as opposed to q24h dosing) would achieve greater inhibition of ACE with the same q24h total dose (20). Furthermore, a different study of single dose enalapril and benazepril at a dosage of 0.5 mg/kg indicated a much shorter duration of effect, with ACE suppression lasting < 12 hours (21), and a recent retrospective study in dogs with valvular heart disease suggested improved outcomes with q12h dosage (22).

There are several reasons why developing consistent recommendations for dosing of ACEi has proven challenging in veterinary medicine. Historically, ACE activity was used as a surrogate for RAAS activity. Recently, however, ACE activity has been shown to be an inefficient measure of RAAS activation. Numerous studies in humans and dogs have shown a lack of correlation between circulating ACE activity and Ang II concentrations (4,23). A second challenge in developing scheduling recommendations is the significant chronobiological modulation of the RAAS. Previous experimental models of RAAS activation failed to consider the chronobiology of the RAAS, while contemporary research has shown that biomarkers of the renin pathway are subject to circadian variations in dogs (4,23,24). Finally, existing PKPD studies on the effect of various ACEi have not consistently sampled biomarkers of *alternative* RAAS activation in addition to biomarkers of *classical* RAAS activation.

Overall, although the effects of ACEi, such as benazepril, on ACE activity have been fairly well characterized, and the benefit of ACE inhibition in CHF has been definitively established in several clinical trials in both humans and dogs (0.25 to 1.0 mg/kg q12h-q24h), little is known about the effect of benazepril on the alternative RAAS pathway in either species. Understanding the dose-dependent effects of benazepril on biomarkers of both the *classical* and *alternative* RAAS pathways in dogs would allow exploration of benazepril dosages that produce a downregulation of the *classical* RAAS while preserving, or upregulating, the *alternative* RAAS. This would translate into an optimization of the clinical benefit. Accumulating data that inform such a nuanced approach to dose optimization in dogs would provide valuable translational information for similar dose optimization of ACEi in humans. To model and predict the dose-dependent effects of benazepril on the classical and alternative arm of the RAAS, we aimed to build a nonlinear mixed-effects (NLME) model of benazepril PKPD. NLME modeling of benazepril PKPD had already previously been shown to be an efficient method for describing the effect of benazepril on the classical RAAS in canines and is a well-accepted framework for building PKPD models (11).

To produce data for this modeling and simulation effort, nine healthy beagles were intensively sampled while administered benazepril at various dosages and frequencies. After producing the data, our objective was to use a quantitative-systems pharmacology (QSP) model to characterize the PKPD relationship of benazepril(at) on biomarkers of the RAAS which are relevant to CHF pathophysiology and associated with morbidity/mortality {angiotensins I, II, III, IV, (1-7)}. QSP modeling is a subgroup of PKPD models which seeks to describe the behavior of a pharmaceutical in terms of the biology of its mechanism of action. After developing and calibrating the model, we further developed a software implementation of the benazeprilat-RAAS QSP model, which is capable of rapidly simulating the effect of benazepril HCL at various doses in a larger population of virtual dogs. By developing an easy-to-use simulation interface for our model, the objective of this work was to make a first prediction of the optimal dose/time of benazepril administration in dogs in support of future investigations in patients with CHF.

## Results

### Animal Safety

All study dogs received all oral doses of benazepril as intended. Dogs were monitored for adverse effects associated with benazepril labeling as well those associated with general animal welfare e.g. vomiting, diarrhea, inappetence, weakness/hypotension, fatigue, incordination, hypercreatininemia, etc.. No adverse effects were observed in the animals during the course of the study, and serial complete blood counts and chemistry panels performed showed no evidence of hematologic or biochemical abnormalities from benazepril dosing.

### Data Mining

Data were collated and standardized for mathematical modeling as instructed in the Monolix documentation (38). Except for standardization of units as molar amounts and concentrations, raw data were left untransformed. Doses were transformed using the molecular weight of benazepril HCl, while concentrations were transformed using the molecular weight of the active metabolite – benazeprilat. Data were reviewed for bulk trends and data quality both before, during, and after mathematical modeling.

Data below the lower limit of quantification (LLOQ) were modeled by adding to the likelihood function a term describing the probability that the true observation lies between zero and the LLOQ, which is equivalent to the M3 method implemented in the NONMEM (Non-linear Mixed Effects Modeling) software.

The log_10_ time-course of benazeprilat as well as the relevant RAAS biomarkers are reproduced in **Fig. 2**. Of note, there was some background experimental noise in pharmacodynamics of some biomarkers that ultimately reduced model prediction quality. The noise was most prominent in the biomarker Angiotensin III (2-8) (i.e. AngIII), where the order of the limit of quantification (2.5 pmol/L) was approximately half of the measurement at the 3^rd^ quartile (5.1 pmol/L). Suspected outliers were flagged and tested as model covariates for statistical significance. However, none of the flagged data points were determined to be significant enough outliers to exclude from model building.

**Figure 2:**
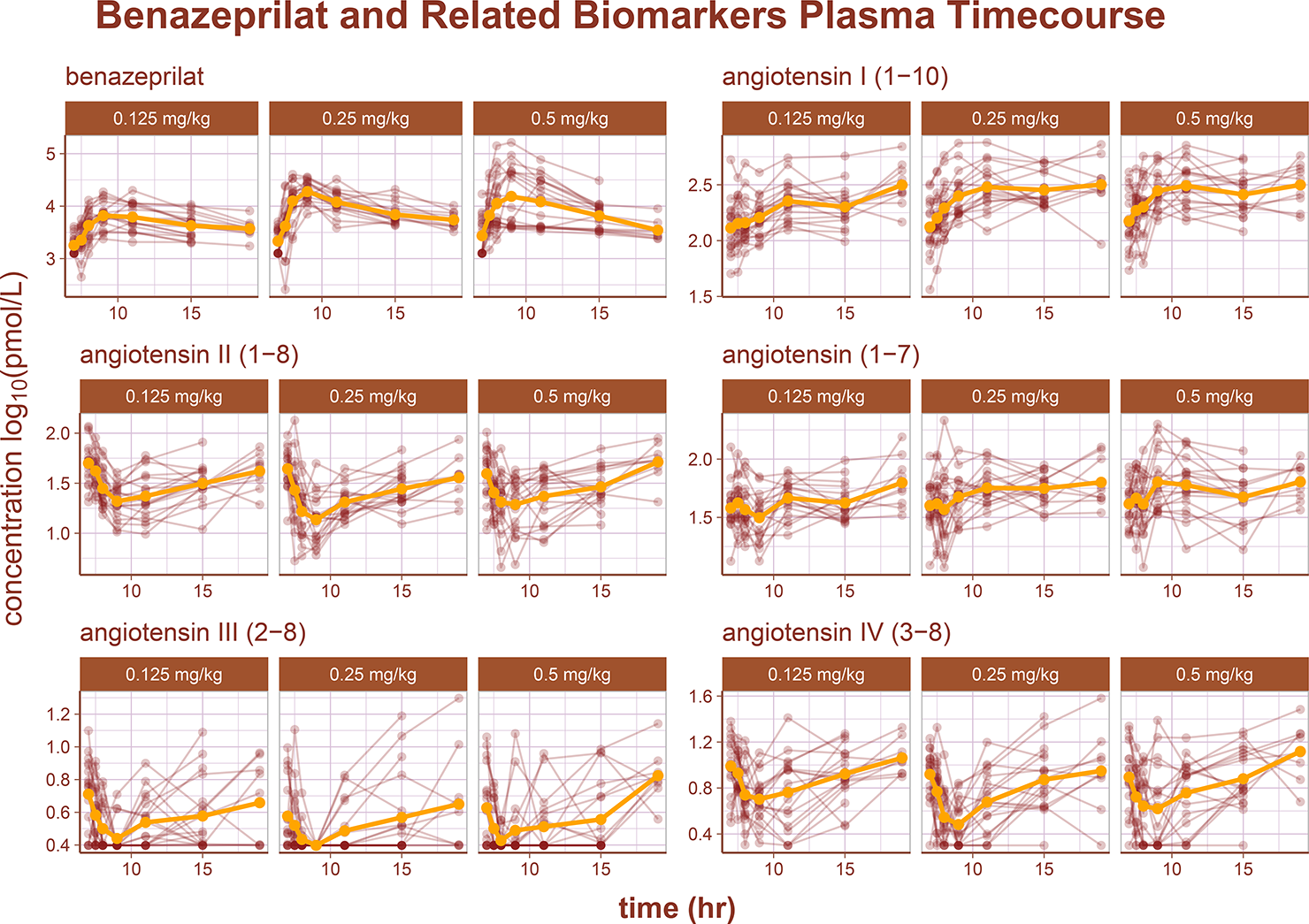
RAAS Biomarkers Timecourse. An overview of the plasma timecourse of several RAAS biomarkers as well as benazepril’s active metabolite, benazeprilat. Each subject’s timecourse is indicated with a red line and points. The golden curve is the mean timecourse value.

Following is a summary of the model building process. The base empirical version of the full model was largely an adaption of the benazeprilat PKPD model reported in Mochel et. al (23). In all, over 100 different structural modifications were tested, starting from the empirical base model, to produce our final QSP model. To simplify results reporting, the most important modifications tested are summarized in the following two sections. Despite the division of sections into PK and PD, after building a base model to work from, all model fits were performed on the full PKPD dataset.

### PK Model Building

The PK portion of the base model was a conventional 2-compartment mammillary model with saturable exchange between the central and peripheral compartments. Building on this initial model, several modifications of the PK structure were evaluated. First, several standard compartment variations were tested i.e., using 1-, 2-, or 3-compartment disposition functions. Overall, a 2-compartment PK model outperformed the other candidate models based on the precision of individual parameters and overall quality of fit. Second, *non-specific* (low affinity, high capacity) binding of benazeprilat to plasma proteins was represented by a 3^rd^ compartment *within* the central compartment, i.e., representing the free circulation of benazeprilat. The volume of the non-specific binding compartment (*V_ns_*) is a representation of the relative binding capacity of benazeprilat which is distributed in plasma but is not freely circulating or interacting with ACE. Therefore, the total amount of measurable benazeprilat in plasma is a combination of the amount non-specifically bound to plasma proteins (*I_ns_*) (low affinity, high capacity), the amount specifically bound to ACE (high affinity, low capacity) and the amount of benazeprilat in free circulation (*I_free_*) (4,20). The variable *I* was chosen to represent benazeprilat as it *<u>i</u>nhibits* ACE activity.

Zero-, first-, mixed-, and sequential absorption structures were tested to model drug absorption from the depot compartment (i.e., intestinal lumen). A model largely equivalent to sequential absorption, but made to be continuous, was found to outperform other competitive models. This model substructure uses a series of 1-order absorptions but can be seen as a continuous analog to a sequential 0-/1-order absorption.

In this model, the first depot compartment for benazepril after oral administration was called *1abs*. First-order absorption either occurred immediately at rate *ka_1_* to the compartment of freely circulating benazeprilat (*fr*), or absorption was delayed by an absorption rate *ka* through an intercompartment that was pre-circulatory (*pr*). As is typical, quantity of benazeprilat passed between compartments is called *I_m, n_*, for inhibitor, where indexes m and n represent origin and destination compartments, respectively. *F_bio_* represents the total bioavailability (**Eq. 4**). Doses are administered in benazepril hcl, but are measured as the metabolite – benazeprilat. To reduce complexity in modeling, but preserve absorption and benazepril to benazeprilat conversion variance, all bioavailable benazepril is treated as benazeprilat in the model. *F_bio_*, or total bioavailability, is estimated in the model purely to preserve this variance and to reduce numerical instability in estimation. However, without IV data, the final estimated *F_bio_* does not have a firm pharmacological interpretation.

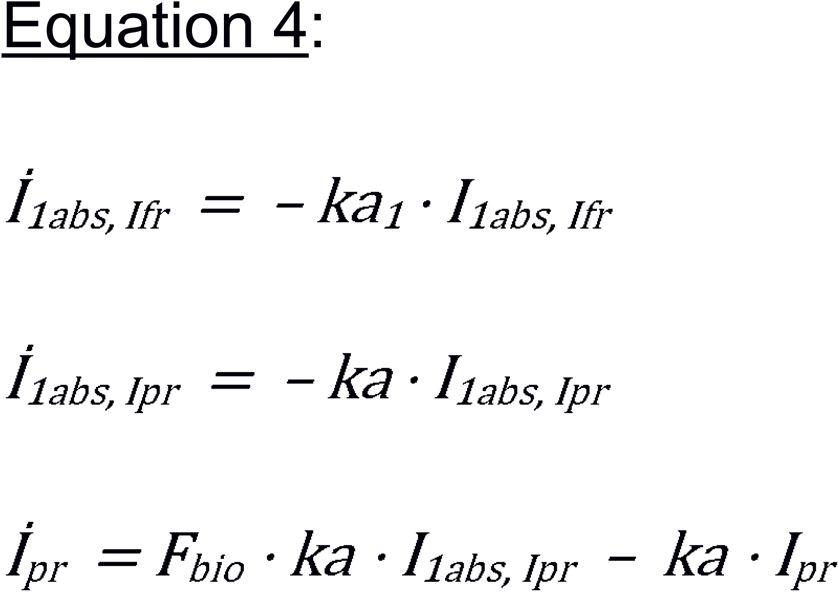

In summary, the final mammillary model *without ACE binding kinetics* (**Eq. 5**) was a 2-compartment PK model with nonspecific protein binding represented by a 3^rd^ compartment (*I_ns_*), and a continuous analog to sequential 0-/1-order absorption from some depot compartment.

Rate of exchange between compartments were governed by rates *k_f, g_* where *f* and *g* (*f ≠ g*) were each one of either free circulation (*fr*), tissue (*ts*), or non-specifically bound in circulation (*ns*). Residual error was best modeled using a normal-proportional error function (**Eq. 9**). The only exception were rates of elimination which were written as *k_Cl, d_* where clearance represented that the parameter was derived from clearance and *d* was the compartment of origin.

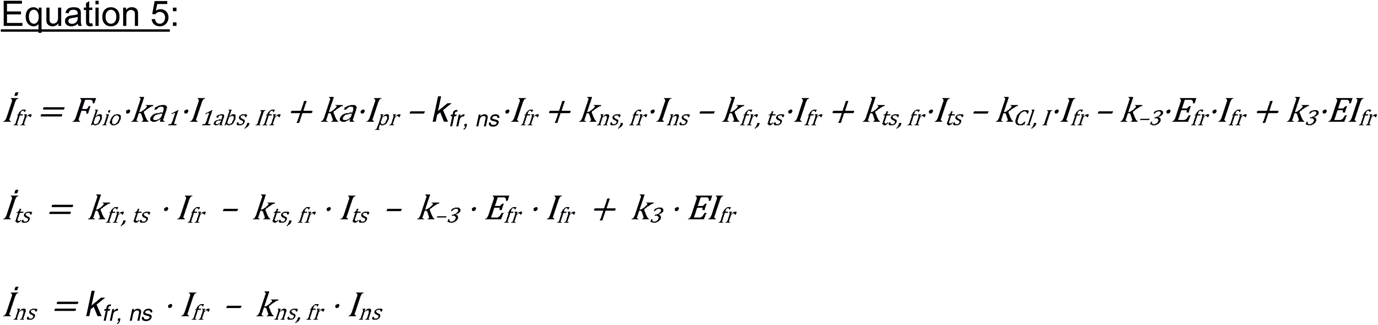

### PKPD Model Building

Benazeprilat primary mechanism of action is inhibiting ACE to prevent the catalysis of AngI into AngII. To account for this mechanism, a logistic saturation model was first implemented. However, the superior model for predicting benazeprilat ACE inhibition was found to be the differential Michaelis-Menten model of catalysis inhibition with ACE being the *<u>e</u>nzyme* (*E*), benazeprilat the *<u>i</u>nhibitor* (*I*), AngI being the *<u>s</u>ubstrate* (*S*) and AngII being the *<u>p</u>roduct* (*P*) (**Eq. 6**). The distribution of ACE across tissue (*ts*) and free circulation (*fr*) was also considered. The nomenclature used throughout Eq. 6 is consistent with previous descriptions of the Michaelis-Menten model (39).

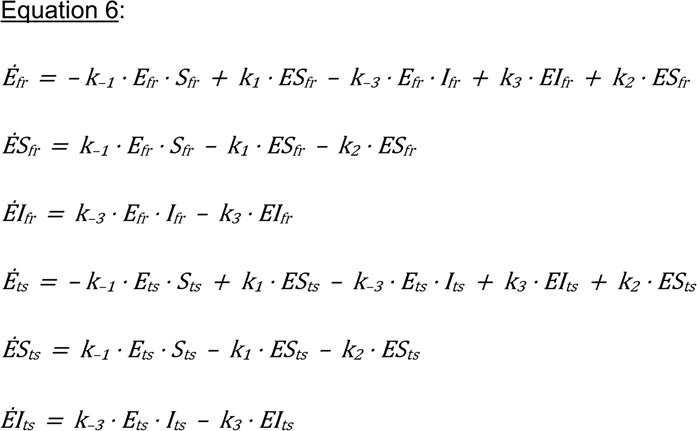

Two-compartment mammillary models governed the kinetics of biomarkers AngI, AngII, and Ang(1-7). The amount in these compartments were respectively represented by *S* (*<u>s</u>ubstrate*), *P* (*<u>p</u>roduct*), and *Ang(1-7)*. The two compartments for these angiotensins were called free circulation (*fr*) and tissue (*ts*). Conversion steps in the *classical* and *alternative* RAAS pathways were modeled through a series of catalytic steps, as previously described (40).

At last, the function *f_CT_(t)* governs the effect of chronobiology on the production rate of the substrate (*r_s_*). *f_ct_(t)* is a scaled cosine function where the wavelength (or period) is matched to 24 hours, the relative maximum amplitude is the scalar *PRA* (peak renin amplitude), and the scale of that amplitude is governed by *δ_24hr_*. Chronobiology is herein only modeled relative to AngI production (**Eq. 7**).

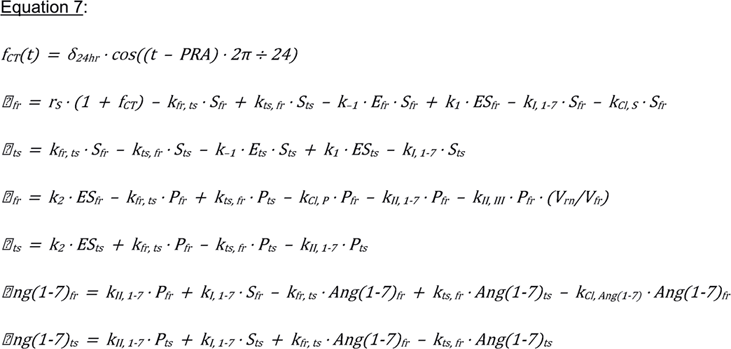

The catalyses of AngII to AngIII, and AngIII to AngIV were modeled via a series of catalytic conversion models (**Eq. 8)**. Cleavages of AngII to AngIII, and AngIII to IV, are primarily performed by renally-bound aminopeptidases A and N, respectively (41–43). *V_free_* was subdivided into two circulatory system volumes of distribution; a small renal volume (*Vrn*) and a larger plasma volume (*Vpl).* All catabolism of AngII to AngIII, and AngIII to AngIV were linked to the renal volume as this is where aminopeptidases A and N are physiologically located.

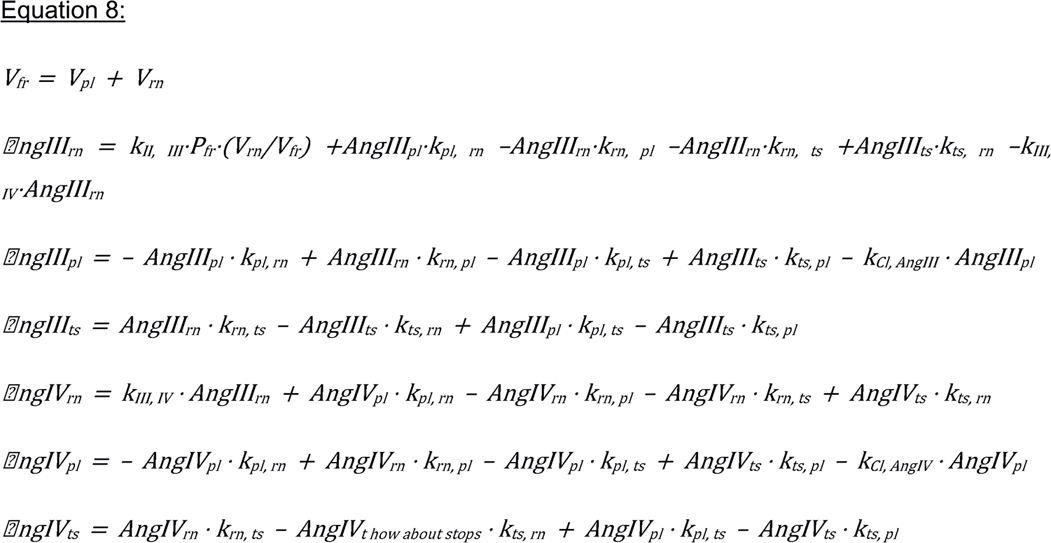

All analytes residuals were best described by proportional error models (**Eq. 9**), with the concentration of a given biomarker scaled by *ε*. *ε* is a normal distribution distributed with standard deviation b, i.e. *ε ~ N(0, b)*.

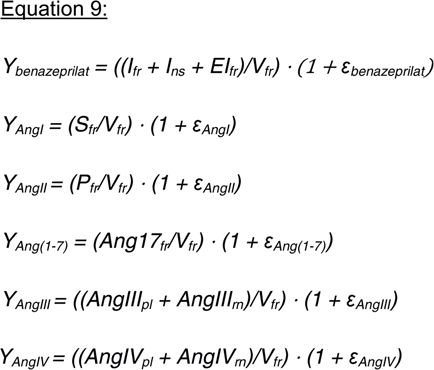

ANOVA tests on covariates indicated that model performance would not be significantly improved by the inclusion of any covariate effects. The full model written in Mlxtran is available in the supplemental files, and a model diagram detailing the full structure is reproduced in **Fig. 3**. In **S1 Table**, the reader can find a detailed description of all mathematical symbols defined in the results section.

**Figure 3:**
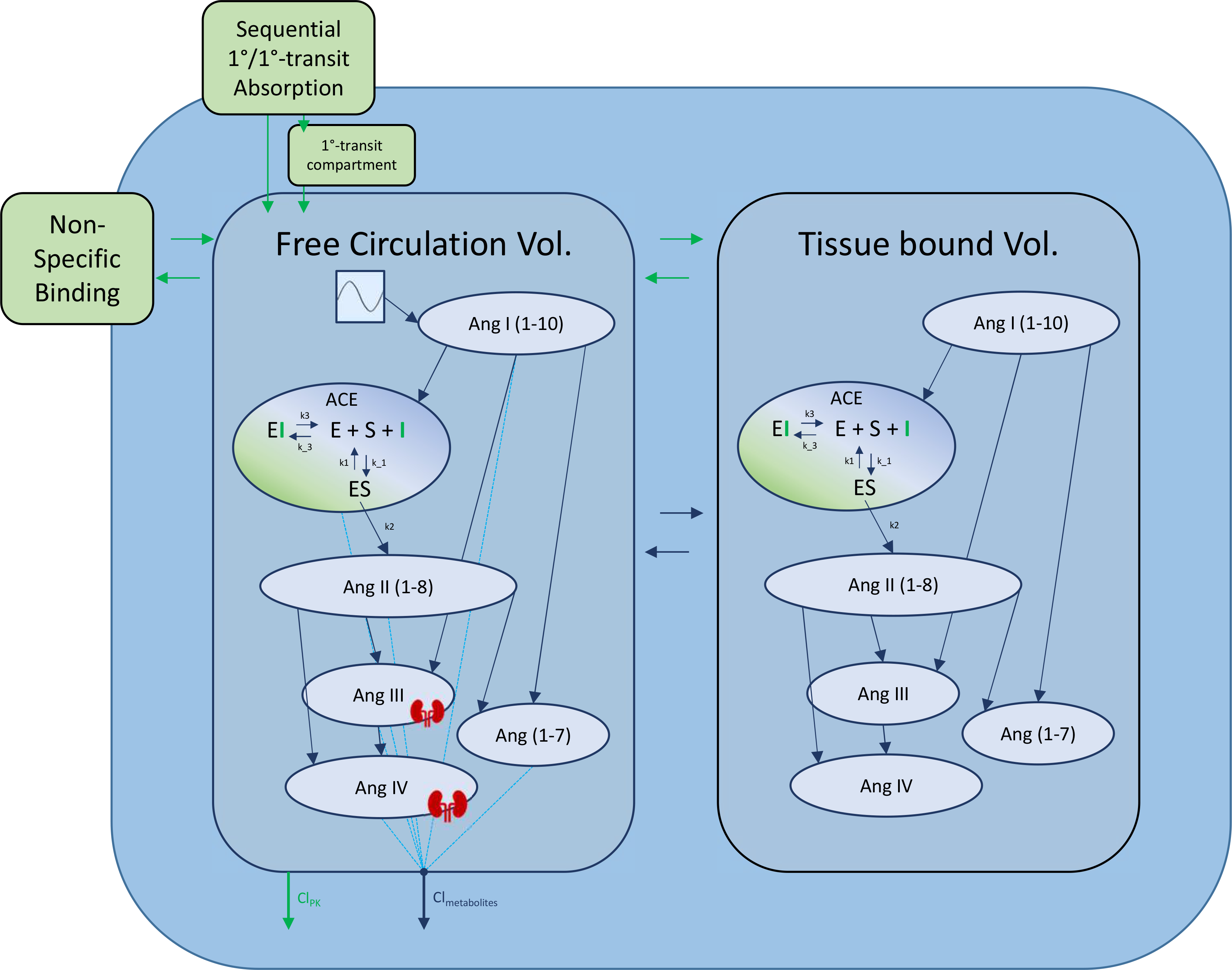
Detailed Model Diagram. Detailed diagram of the final model structure. Benazeprilat pharmacokinetics were modeled using a ***2-compartment model*** with a ***mix of 1-order and 1-order delayed by 1-order transfer absorption*** from the depot compartment. Both volumes of distribution, *free* and *tissue*, were modeled with a fixed amount of ACE with which Benazeprilat could act on. ***Non-specific binding*** affected the free circulation compartment. A series of ***direct response models*** were used to describe the transformation of angiotensin I into its various metabolites. The free volume of distribution was subdivided into ***plasma*** and ***kidney*** volumes for Ang III and Ang IV. A Michaelis-Menten kinetic model of inhibitor, substrate, and enzyme interaction was used to describe the competitive inhibition of ACE by benazeprilat. **k_3_** and **k_−3_** were the parameters governing ACE-benazeprilat (enzyme-inhibitor) association and dissociation, while **k_1_** and **k_−1_** determined the rate of ACE-angiotensin (enzyme-substrate) association and dissociation. **k_2_** controlled the production rate of angiotensin II from angiotensin I via ACE. An **independent clearance** for each metabolite as well as benazeprilat controlled the rate of removal of various molecules from the plasma.

### Model Fit Evaluation

Inspection of the SAEM search and a sensitivity analysis on initial parameter values revealed a stable and precise search for all parameter estimates. The final selected model had high precision in parameter estimates as evaluated via RSE (majority of estimates <35%). A summary of model parameter estimates, including typical value, RSE (%) and inter-individual variability (IIV) can be found in **Table 1**.

**Table 1:**
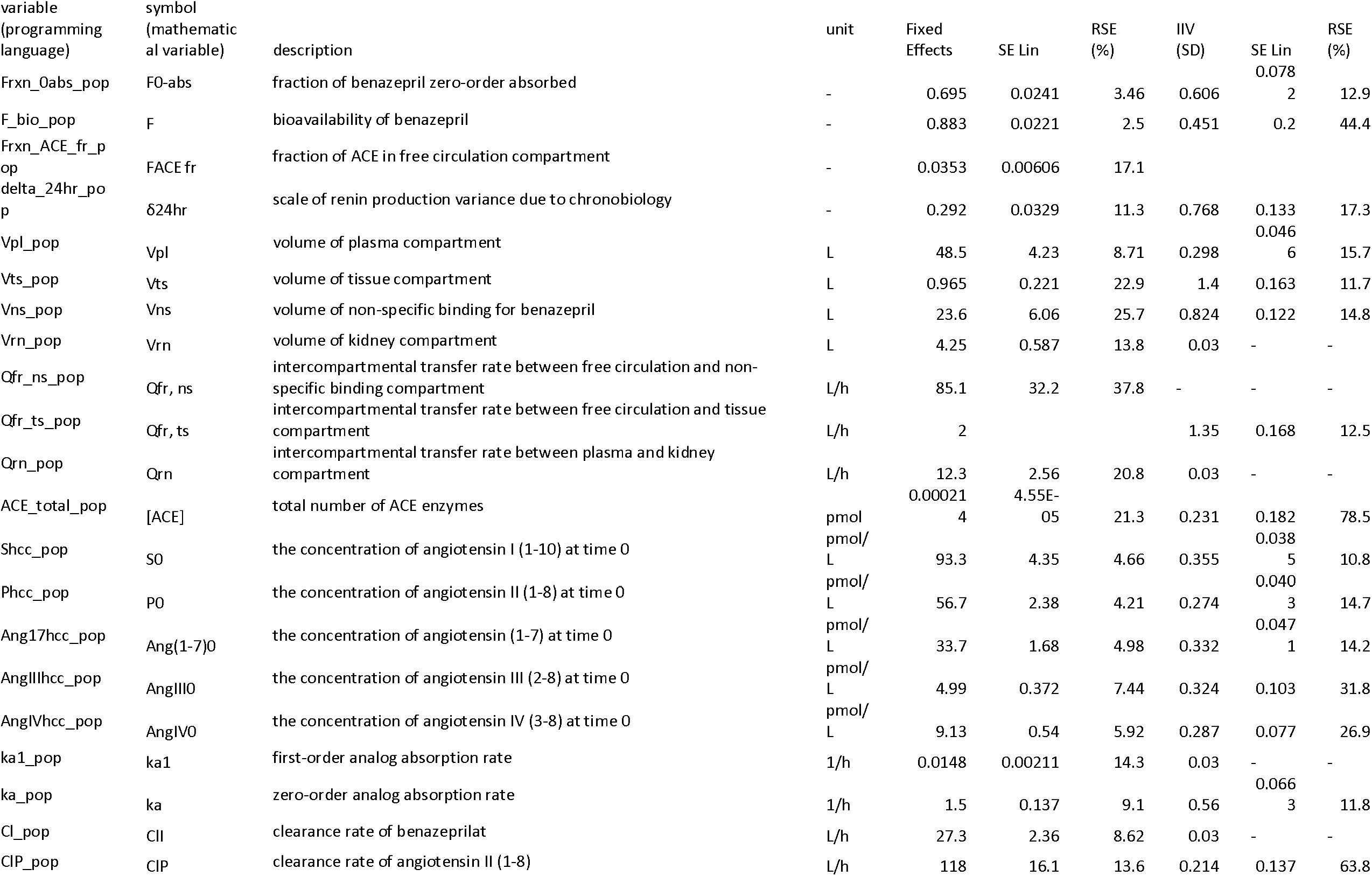
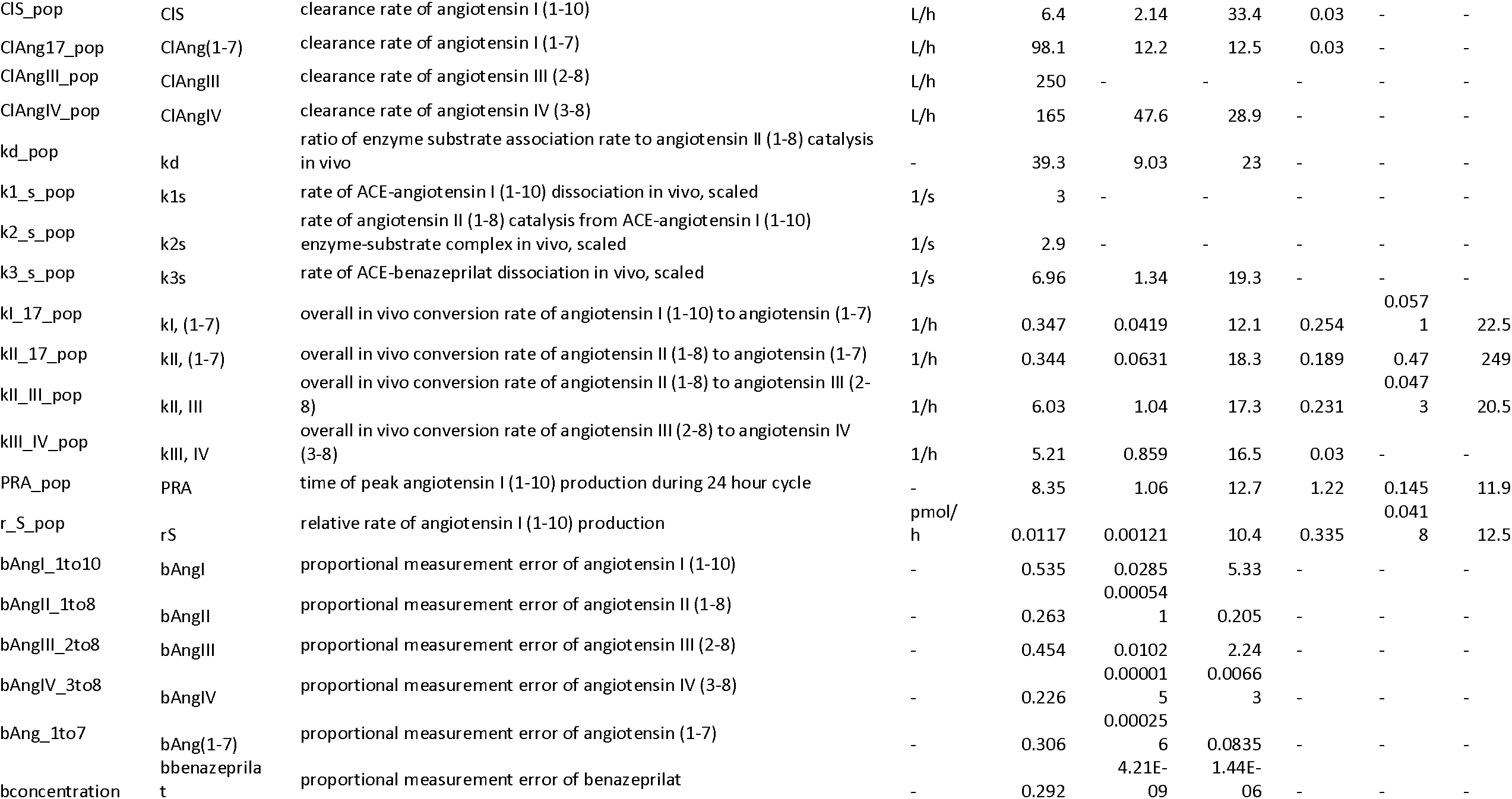
Parameter Estimates. The full table of parameter estimates. In the first column, the name of the parameter used in the computer code is given. Then, the variable name for the mathematical equations is given. A description of the parameter and units are given in the next two columns. Finally, the actual value of the estimate along with error estimate, gamma (standard deviation) estimate, and the error of the standard deviation estimate. Low residual error estimates on the table of parameters indicate that the model is not over-parameterized and the identification of parameters is precise.

Inspection of goodness-of-fit summary plots (**Fig. 3–5**) indicate that benazeprilat predictions from the model are largely in line with experimental measurements. Importantly, the final PKPD model, which enabled the simultaneous fit of all angiotensins, was found to characterize the time-varying changes of the both the classical and alternative arm of the RAAS satisfactorily, as shown by the standard goodness-of-fit diagnostics of observations vs predictions (**Fig. 3**), the individual predictions (**Fig. 4**), and the simulation-based validation diagnostics (i.e., NPDEs, **Fig. 5**).

**Figure 4:**
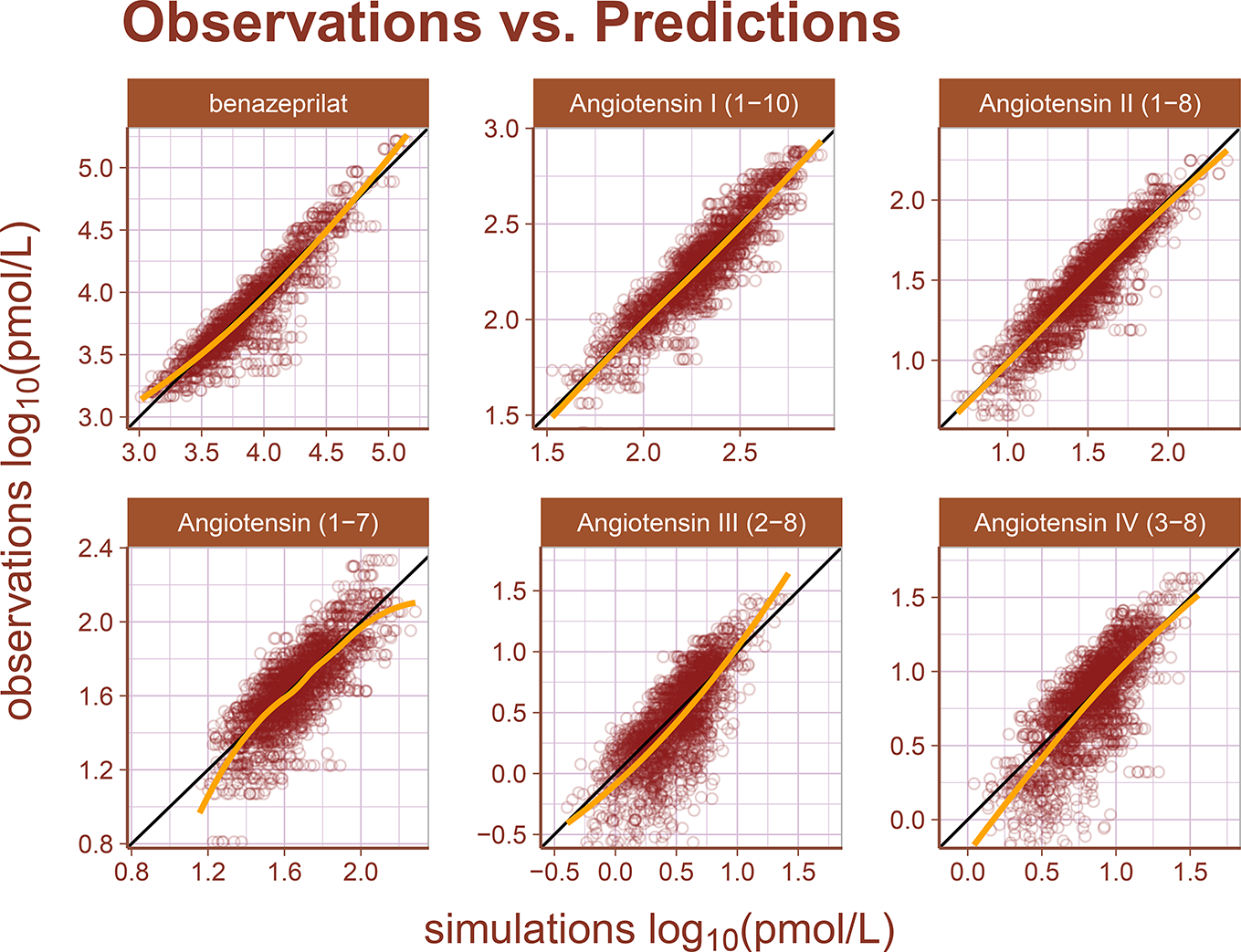
Observations vs Predictions. The observations plotted against the predictions for all metabolites and drug concentration data. This gives a complete picture of model performance. The golden line is the LOESS curve showing the correlation between observations and predictions. The black line plotted diagonally represents an ideal model performance with no misspecification. The general agreement between LOESS and idealized curve indicate that there is little misspecification in model structure.

**Figure 5:**
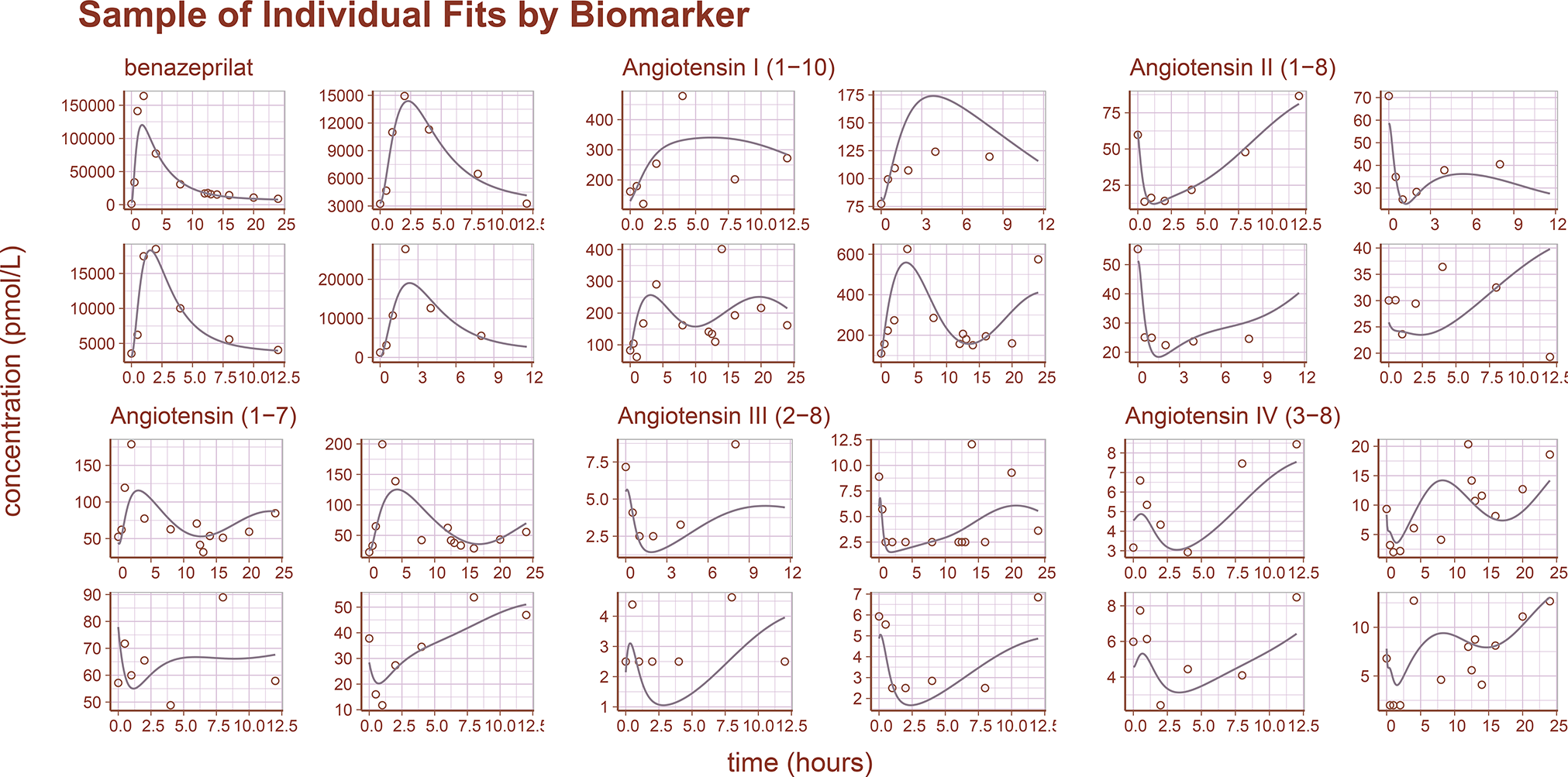
Sample of Individual Predictions. A sample of individual observations vs predictions randomly sampled from the concentration and metabolite data. The general agreement between plasma concentration timecourse and individual predictions indicates that the model reproduces the observations with high accuracy.

### Simulation Engine

There are three primary views in this application (**Fig. 6**). In all views, the time of first dose of benazepril is specified in a 24-hour clock format. On the left-hand side of the application is a menu for specifying dosing, parameters of the simulation, and modalities for calculating the area under the effect curve (AUEC) that quantifies the effect of the active benazeprilat on the RAAS at various doses vs. placebo control. Note that the application menu can be hidden to increase the size of the plotting area.

**Figure 6:**
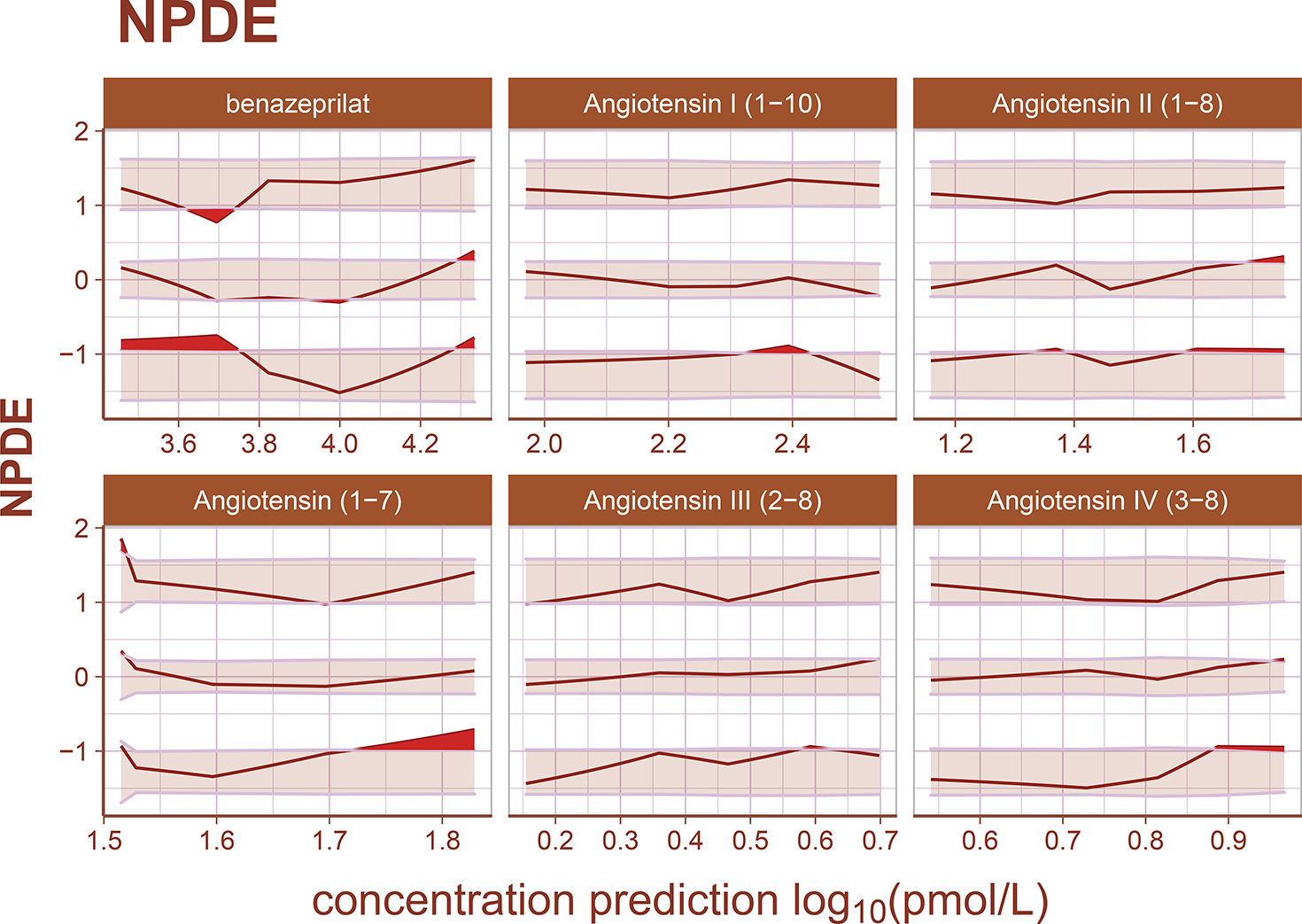
Normalized Prediction Distribution Errors. Normalized prediction distribution errors (NPDEs) are an analog to residuals used for diagnosing both model structural misspecifications as well as the performance of the residual error model. The distribution of a well-specified model is normal, ideally. Bands represent the 90% prediction band for the 95^th^, 50^th^, and 5^th^ percentiles, respectively. Curves are the observed percentiles for the 95^th^, 50^th^, and 5^th^ percentiles, respectively. Data are binned at regular intervals to derive these average trends. For each binning range, if the structural model fits the data well, observed percentiles will be symmetrically distributed across a 50^th^ percentile curve which falls within the 50^th^ percentile band. If the error model is well specified, observed percentiles will fall within prediction bands. Any misspecification is ideally random. The model appears to underpredict angiotensin (1-7) for small measured values slightly, but otherwise there is high agreement between model and data.

#### Application Menu

The left-hand menu is split into three tabs which allow the user to define parameters of the simulation. The *dosing* tab permits the user to define the dosing schedule in terms of time of first dose, number of doses, size of dosage, and interdose interval. The *simulation* parameters tab gives the user access to the timescale of the simulation, the fineness of the grid used for simulation, and the sample size used to calculate the median and prediction intervals of the simulated PKPD. Finally, the *AUEC* tab provides a means to compare pharmacodynamic effects between competing dosing scenarios by defining a time period for which to calculate AUEC estimates.

#### Prediction Distribution View

The first tab gives the user tools for analyzing the distribution of responses after a single schedule of benazepril. The distribution is specified in terms of median effect (blue line), median effect of placebo using same simulated individuals (dashed black line), and 90% prediction interval (blue bands) in steps of 10% i.e., 5% to 95%, 15% to 85%, etc. The AUEC of treatment vs. placebo can be compared for the timespan between the dashed vertical lines. The percent difference between those two AUECs is documented in the hovering label.

#### Dosage Comparison View

The second panel allows the user to compare up to four competitive dosing schedules to placebo. In this panel, the user can see the median response (key at bottom) and the placebo effect (dashed black line), but not the distribution of responses. On the right-hand side of the page, the user can compare the percent difference from placebo in the RAAS components modeled in this study by paging through the various data tables. These comparisons are percentages relative to placebo.

#### Documentation View

The last view is simply documentation on the model that powers the simulation engine and general recommendations for using the software. It also provides a brief summary of the design of the application and gives a full reproduction of the R code that makes up the model. In this view, the application menu also gives a brief summary of user warnings.

#### Dosage Comparisons

As a final consideration in our study, we directly compared four dosage scenarios in our simulation engine: 0.25 mg/kg q24h in the AM, 0.25 mg/kg q24h in the PM, 0.25 mg/kg q12h, and 0.5 mg/kg q12h. In our simulation application, we set the engine to compare the median AUEC of 500 dogs (matched between virtual trials), at a sampling rate of 500 times over a period of 25 virtual days. The median AUEC comparison was made on day 20 over a period of 24 hours. The long virtual time of simulation assured the simulated dogs reached steady state PD of benazeprilat. For 0.25 mg/kg q24h, we saw an approximate 55% decrease compared to placebo for AngII and a 94% increase in Ang(1-7). With a schedule of 0.5 mg/kg q12h, we saw an approximate 80% decrease versus placebo for AngII and 135% increase in Ang(1-7). We saw a greater daily biomarker variance in SID dosing. *Summary results* are tabulated in **Table 2** while median time-courses are plotted in **Fig. 8**.

**Table 2:** Simulation Summary. Four administration schedules are compared in this simulation scenario. 1). 0.25 mg/kg every 24 hours at 8am 2). 0.25 mg/kg every 24 hours at 8pm 3). 0.25 mg/kg twice a day at 8am and 8pm 4). 0.5 mg/kg twice a day at 8am and 8pm 500 individuals were simulated for each scenario (for a total of 2500 individuals, with placebo). Curves are the median timecourse of the molecule from this simulated population. Comparisons between schedules are made by calculating the percent difference between the median AUC and placebo (of each schedule). Median timecourses are plotted in **Figure 8**. There was little difference between morning and night dosing at steady-state. Angiotensin (1-7) production and angiotensin II reduction were approximately doubled (vs. placebo) by increasing to twice a day dosing. The effect of increasing the dosage was more marginal, but still significant (approximately 15%).

| schedule<br>(mg per<br>kg) | Ang<br>(1-7)<br>(%) | 5<br>%ile | 95<br>%ile | AngI<br>(1-10)<br>(%) | 5<br>%ile | 95<br>%ile | AngII<br>(1-8)<br>(%) | 5<br>%ile | 95<br>%ile | AngIII<br>(%) | 5<br>%ile | 95<br>%ile | AngIV<br>(%) | 5<br>%ile | 95<br>%ile |
| --- | --- | --- | --- | --- | --- | --- | --- | --- | --- | --- | --- | --- | --- | --- | --- |
| q24h 0.25 | 94.8 | 6.8 | 284 | 155 | 34.4 | 417 | -55.6 | -78.9 | -8.2 | -56.4 | -85 | 12.8 | -56.2 | -85.2 | 10.7 |
| q24h 0.25 | 94.1 | 5.7 | 282 | 155 | 31.7 | 411 | -54.7 | -79.2 | -6.1 | -56.2 | -85.3 | 14.9 | -56.1 | -85 | 11.4 |
| q12h 0.25 | 120 | 21.6 | 324 | 196 | 50.6 | 504 | -69.7 | -86.6 | -32.7 | -70.8 | -90.7 | -17.3 | -70.7 | -90.6 | -18.5 |
| q12h 0.5 | 135 | 31 | 342 | 222 | 60.7 | 559 | -79.6 | -91.8 | -45.7 | -80.1 | -94.3 | -36.9 | -80.8 | -94.3 | -39.7 |

**Figure 7:**
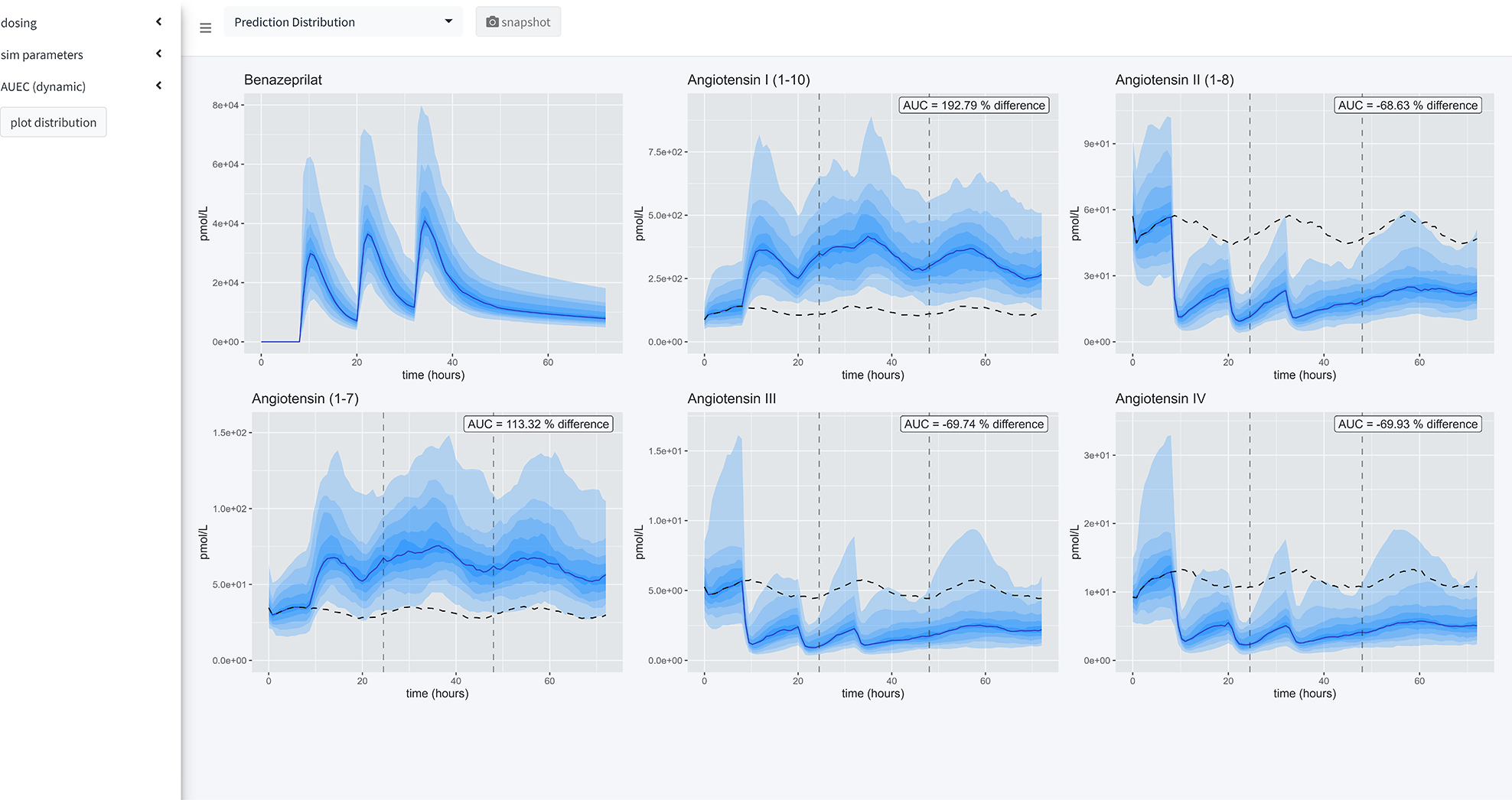
Application Views. Our application provides a user-friendly way to apply our model of RAAS response to various administration schedules of benazepril. A collapsible left-hand widget allows the user to specify the simulation. The application has 3 separate panels. In the first panel, a single dosage scheme can be applied to a large simulated population of animals. Then the <u>prediction distribution</u> of simulated patient responses is plotted for study. In the second panel, the user can make a <u>comparison</u> between several proposed administration schedules. The plots produced in this panel depict the median timecourse of the various metabolites in response to the proposed schedules. The user also has access to an x-axis zoom and AUC comparison summaries on the right-hand side. The final panel is simply <u>documentation</u> of the simulation engine code and tips for usage.

**Figure 8:**
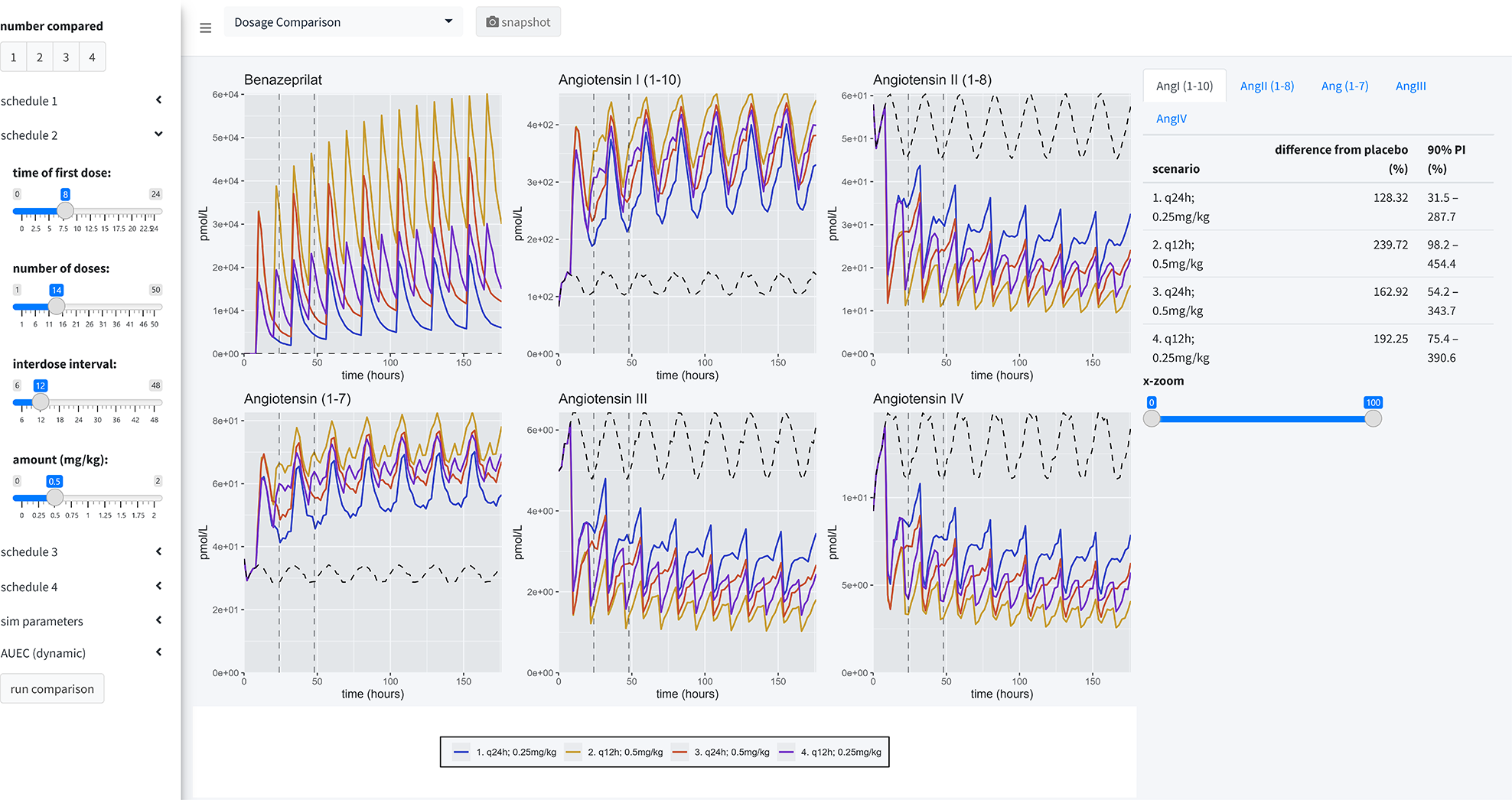
Simulation Summary. Four administration schedules are compared in this simulation scenario. 1). 0.25 mg/kg every 24 hours at 8am 2). 0.25 mg/kg every 24 hours at 8pm 3). 0.25 mg/kg twice a day at 8am and 8pm 4). 0.5 mg/kg twice a day at 8am and 8pm 500 individuals were simulated for each scenario (for a total of 2500 individuals, with placebo). Curves are the median timecourse of the molecule from this simulated population. Comparisons between schedules are made by calculating the percent difference between the median AUC and placebo (of each schedule). Summary results are tabulated in **table 2**. There was little difference between morning and night dosing at steady-state. Angiotensin (1-7) production and angiotensin II reduction were approximately doubled (vs. placebo) by increasing to twice a day dosing. The effect of increasing the dosage was more marginal, but still significant (approximately 15%).

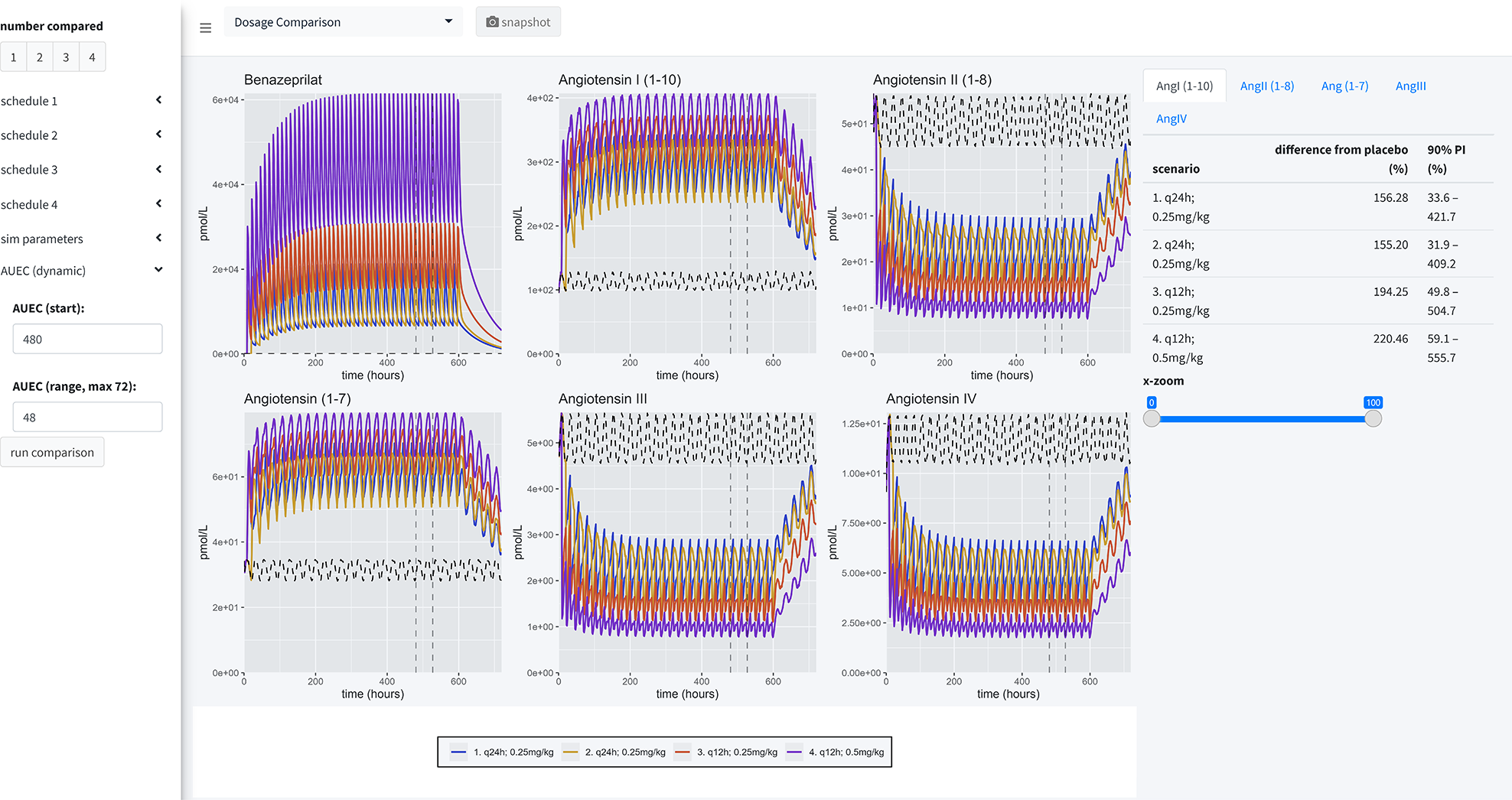

## Discussion

In both canines and humans, *classical* RAAS overactivation plays a key role in the pathogenesis and development of CHF (4). To modulate *classical* RAAS overactivation in CHF, there is a substantial history of using ACEi such as benazepril in both species (2,3,44). Improving our understanding of the effect of ACEi on CHF in canines has the potential to benefit therapeutic management of CHF in humans and *vice versa* (4). Activation of the *alternative* RAAS pathway is characterized by catalysis of AngII to Ang(1-7) by the enzyme ACE2. In turn, Ang(1-7) activates Mas receptors (Esteban PloS One 2009). In direct contrast to the effect of *classical* pathway overactivation on CHF, activation of the *alternative* pathway is associated with improved clinical outcomes via reduced and preserved ejection fraction and reduced risk of heart failure.

An ideal therapeutic for CHF would modulate both the *classical* and *alternative* RAAS pathways, downregulating the activity of the *classical* RAAS pathway while preserving or upregulating the *alternative* RAAS pathway (13). Although the effect of ACEi on ACE activity and AngII has been fairly well-characterized both in the veterinary and human literature, little is known about the effect of ACEi on the *alternative* arm of the RAAS. Therefore, any further dose optimization of ACEi is dependent on studying the effect of this therapeutic drug on the *alternative* pathway. Characterizing this effect in dogs essentially furthers our understanding of how to optimize therapeutic management of canine CHF while providing valuable insight into the molecular effects of ACEi for translation to human CHF (45).

Additionally, in humans and in dogs, the recommended dosage range for benazepril is quite broad and there is no clear consensus on the ideal dose in patients with CHF. PKPD studies comparing various doses of benazepril in healthy dogs have not provided consistent recommendations. In King et al. (1995) a single PO dose of benazepril effectively suppressed ACE activity for up to 24 hours, and that ACE inhibition in plasma was independent of dosage ≥ 0.25 mg/kg (17). Subsequent reanalysis of these data using PK modeling suggested that q12h dosing would achieve greater inhibition of ACE with the same total q24h dose (20). Later, Hamlin & Nakayama (1998) found a single dose of benazepril at 0.5 mg/kg suppressed ACE for < 12 hours (21). Lastly, a recent retrospective study in dogs with valvular heart disease suggested improved outcomes with q12h dosage (22).

In this study, we have attempted to address knowledge gaps in optimal benazepril dosing by describing the dose-dependent effects of benazepril on biomarkers of both the *classical* and *alternative* RAAS pathways in dogs. The solution we implemented to address these gaps in knowledge was to build <u>a modeling and simulation platform of the effect of benazepril on both arms of the RAAS.</u> This simulation engine allows for exploration of benazepril dosages that produce both a substantial downregulation of the *classical* RAAS while preserving or upregulating the *alternative* RAAS. This allows for side-by-side comparison of several dosage schemes using virtual clinical trials and ultimately optimization of clinical benefits.

Several metrics were used in assessing model performance. Population-level goodness-of-fit diagnostic plots like observations vs. predictions and NPDEs indicated that structural misspecification was very low. Focusing on individual predictions, we see a dynamic model capable of fitting complex individual variations without being overly fit to noise and spurious trends in the data. Importantly, for such a large set of parameters, our precision of parameter estimates was very high. Some parameters had to be fixed to exploratory values to achieve that precision, but this is an expected outcome when using enzyme kinematic models. Of note, although the precision of these parameters was high, they should not be overly interpreted without experimental verification.

Using our simulation engine, we chose to compare several reasonable dosing schedules, including 0.25 mg/kg PO q24h and 0.5 mg/kg PO q12h – covering the range of dosing schedules most commonly used in the EU and US. Comparisons between dosing schedules were made based on area under the effect curve of the biomarker response (AUEC) relative to placebo.

Chronobiology played a modest role in the scheduling of benazepril in this study. While using the simulation engine to explore various dosages, evening dosing appears to produce the lowest variance in the classical RAAS, morning dosing appears to produce the lowest variance in AngIII and AngIV PD. Morning and evening administration both produce the same relative AUC improvement over placebo across the biomarker panel.

Of the schedules compared, the most robust downregulation of classical RAAS biomarkers and upregulation of alternative RAAS biomarkers was observed with the 0.5 mg/kg PO q12h dose of benazepril. However, this should be put in context because the improvement of 0.5 mg/kg q12h over 0.25 mg/kg q12h is 26.14%, 9.93%, 14.73%, 9.35%, and 10.1% for AngI, AngII, Ang(1-7), AngIII, and AngIV, respectively. Those respective changes in classical and alternative RAAS biomarker concentrations will need to be linked to clinical outcomes in order to inform a clinical dosage selection.

An interesting finding from our simulations is that there is generally high agreement between q24h and q12h dosing of benazepril, as long as the total dosage per day is kept constant. For example, 0.5 mg/kg q24h and 0.25 mg/kg q12h produce similar pharmacodynamic effects on the RAAS, although q12h dosing led to fewer fluctuations of the angiotensins in plasma compared to q24h dosing.

To the best of the authors’ knowledge, this is the first description of a simulation engine designed to optimize dosages of therapeutic drugs in veterinary medicine. Although experimental validation is necessary for bedside application, the model structure readily lends itself to such validation. Many parameters are directly measurable because they have meaningful pharmacological interpretations. For example, the Michaelis-Menten kinetic parameters which govern ACE inhibition could potentially be measured independently – possibly *in vitro*. Additionally, the kinetics of each individual molecule can be measured separately in IV-PK timecourse studies *in vivo*. Furthermore, by using a combination of *in vitro* studies for defining enzyme kinematic parameters, *in vivo* studies of individual metabolite kinetics, literature values for baseline parameters, and allometric scaling of mammillary compartment parameters, this model can be easily adapted into a bedside tool in both canines *and humans*.

There are several practical limitations in this study worth recognizing to guide future model refinement. First, our results were derived from an experimental model of RAAS activation in response to benazepril, rather than a clinical trial in canine CHF patients. Also, the sample size of the study was quite limited compared to a mature clinical trial design. Lastly, the therapeutic window was not considered when making comparisons between dosing schedules. Specifically, there are high doses of benazepril the user can specify in our simulation engine (max. 2mg/kg q6h) that have not been tested experimentally; although safety up to 1 mg/kg q24h has been established in previous studies (22).

## Conclusion

This extensive QSP model of the RAAS in response to benazepril and the development of that model into a tool for bedside optimization of benazepril for CHF and similar diseases is highly novel. By developing an easy-to-use simulation interface for our model, we are now able to make a first prediction of the optimal dose/time of benazepril administration in dogs in support of future investigations in patients with CHF. Beyond the research presented in this manuscript with our tool, simulation tools can continuously expand the impact of scientific research by being used to test new hypotheses surrounding dose optimization and being improved when new data becomes available to refine parameter estimates. The data we measured in an experimental model of RAAS activation has yet to be linked directly to clinical outcomes in CHF, so there is room to expand the model into disease outcomes with a link function. This will be necessary for ultimate application in dosage selection. The most important extension is to experimentally validate a relevant selection of simulations; this option is currently being explored. This model-based approach is now supporting the design of an upcoming prospective multicenter clinical trial in canine patients with CHF to confirm findings from our simulator and refine our model-based predictions with actual clinical information. This clinical trial will help to confirm if the difference observed in PKPD between different dosages is translated into clinical benefits in dogs with naturally occurring CHF.

## Materials and Methods

### Animals

All study procedures were approved by the Institutional Animal Care and Use Committee at Iowa State University (Protocol #19-344).

Nine purpose-bred beagles (5 castrated males and 4 spayed females), 40-42 months old, weighing 9.0-13.5 kg were randomized based on body weight and sex into three oral dosing groups of benazepril. Systemic and cardiovascular health of all dogs was confirmed prior to the study with physical examination, routine laboratory screening (complete blood count, serum biochemical analysis), blood pressure measurement, and echocardiography.

### Housing Conditions

Study dogs were housed in the Laboratory Animal Resources unit at the Iowa State University College of Veterinary Medicine. Dogs were acclimatized to the facility for >1 month prior to the experiment. Dogs were pair-housed in adjoining pens (approximately 2m^2^ per dog or 4m^2^ per pair) on elevated rubber-coated grated flooring. Housing conditions were standardized with ambient temperatures of 18°C, a 12-hour light cycle (07:00 to 19:00), and access to water *ad libitum*. During intensive sampling days only (D1, D18, and D35), dogs were separated into single housing units and water consumption was quantified every 8 hours for the 24-hour period.

After sample collection on baseline sampling days (D-5, D12, and D29), dogs were offered a low-sodium diet (Hill’s Prescription Diet h/d, 17 mg sodium per 100 kcal) at 23:00 q24h for five days to attain a steady activation of RAAS (4,23,24). After data collection on D2, D35, and D36, dogs began a 10-day wash-out period between cycles during which they were offered their standard diet (Royal Canin Beagle Adult, 110 mg sodium per 100 kcal) q24h at 09:00. Volume of low-sodium diet was calculated so that the dogs received the same caloric intake throughout the study.

### Experimental Procedure

This 35-day prospective study was divided into three periods with three different benazepril dosing groups: (A) 0.125 mg/kg q12hr PO, (B) 0.25 mg/kg q12hr PO, and (C) 0.5 mg/kg q24hr PO. All dogs received all treatments using a partial crossover (ABC/BCA/CAB) design. Dogs were sampled in the same order at each time point and exact time of sampling was recorded. Blood sample collection was divided into baseline sampling days (D-5, D12, and D29), sparse sampling days (days 0, 17, and 34), and intensive sampling days (D1, D18, and D35). Baseline and sparse sampling occurred at 07:00.

On intensive sampling days (D1, D18, and D35), blood was collected starting at 07:00 (0 hour, immediately before oral benazepril dosing) and repeated at + 0.5, 1, 2, 4, 8, 12, 12.5, 13, 14, 16, 20, and 24 hours post-dosing. Benazepril (NELIO^®^ 5 mg chewable tablets, Ceva Sante Animale) was administered on intensive sampling days following the 0-hour blood sampling (all dose groups) and the 12-hour sampling (q12hr dose groups only). The benazepril dose was calculated to the nearest 1.25 mg increment.

Venous blood samples were collected from an external jugular or cephalic vein with a 1 inch, 20-gauge or 22-gauge needle attached to a 6 mL syringe. Dogs were kept and maintained in the same position (seated with neck extended) during blood collection. On intensive sampling days, approximately 4 mL of whole blood was collected at each time point with 2 mL transferred to an additive-free collection tube and 2 mL transferred to a lithium heparin tube containing 11.2 μL of dichlorvos prepared as a 6 mg/mL solution in acetonitrile. On baseline days, approximately 6 mL of whole blood was collected with 2 mL placed in an additive-free tube for RAAS analysis, 2 mL placed in an EDTA tube for complete blood count, and 2 mL placed in an additive-free tube for serum chemistry panel. On sparse sampling days, 2 mL of blood was collected and placed in an additive-free tube. All samples intended for pharmacokinetic or RAAS analysis were centrifuged for 15 minutes, after which plasma or serum was transferred into cryovials that were then stored at −80°C for later analysis. Samples for complete blood counts or serum chemistry panels were analyzed by the Iowa State Clinical Pathology Laboratory.

### Analytical Methods

#### Benazeprilat Pharmacokinetics

Plasma benazeprilat analysis was performed by the Iowa State University Analytical Chemistry Laboratory. Benazeprilat and benazeprilat-d5 analytical standards were purchased from Toronto Research Chemicals (Ontario, Canada). Benazepril and benazepril-d5 analytical standards were purchased from Cayman Chemical (Ann Arbor, Michigan, USA). Benazeprilat and benazeprilat-d5 stock standard solutions were prepared at 0.25 mg/mL in 2:1:1 acetonitrile:water:DMSO. The benazepril and benazepril-d5 stock solutions were prepared at 1 mg/mL in acetonitrile. Control beagle plasma was purchased from Equitech Bio (Kerrville, TX, USA). All solvents used for sample preparation and the chromatography portion of the analytical method were purchased from Fisher Scientific (Waltham MA, USA).

A sample volume of 150 μL was fortified with 15 μL of a benazeprilat-d5 solution at 0.1 ppm. Plasma samples were precipitated with 600 μL of acetonitrile containing 0.5% formic acid and vortexed by hand for several seconds. All samples were centrifuged at 10,000 rpm for 5 minutes. A 600 μL volume of each sample was transferred to a clean 2 mL flip-top tube. All flip-top tubes were placed in the CentriVap Concentrator system (Labconco Corp., Kansas City, MO, USA) and concentrated to dryness. All samples were reconstituted in 100 μL of 50:50 acetonitrile:water and centrifuged at 10,000 rpm for 5 minutes prior to LC-MS/MS analysis. All samples were analyzed using an injection volume of 2 μL.

A Vanquish Flex LC pump interfaced with a TSQ Altis mass spectrometer (Thermo Fisher Scientific, San Jose, CA, USA) were used for the analysis. The source conditions were as follows: spray voltage - 3500 V, sheath gas - 40.6 Arb, auxiliary gas - 23 Arb, sweep gas - 0.4 Arb, ion transfer tube temperature - 325 °C, and vaporizer temperature - 350 °C. The total run time of the method was 3 minutes. The resolution of Q1 and Q3 was 0.7 FWHM. The CID gas was set to 2 mTorr. The chromatographic peak width was 2 sec. and the cycle time was 0.2 sec. The mass spectrometer was operated in positive ion electrospray ionization mode. Data was acquired using a multiple reaction monitoring (MRM) method that selected for the benazeprilat ([M+H]^+^ 397.2) and benazeprilat-d5 ([M+H]^+^ 402.2) precursor ions.

The column used for the analysis was Hypersilgold Aq 50 × 2.1 mm, 1.9 μm (Thermo Fisher Scientific, Waltham, MA, USA). Mobile Phase A was water + 0.1% formic acid and Mobile Phase B was acetonitrile + 0.1% formic acid. The column oven temperature was set to 35 °C. The chromatography gradient was as follows: Start at 0% B and linear ramp to 100 %B in 2.0 min, hold at 100% B for 0.4 min, drop to 0% B in 0.01 min, and hold at 0% B for 0.59 min. The flow rate of the method was 0.4 mL/min.

#### Benazeprilat Pharmacodynamics: RAAS Fingerprint

Equilibrium concentrations of Ang I, Ang II, Ang III, Ang IV, Ang 1-9, Ang 1-7, and Ang 1-5 were quantified in serum samples by liquid chromatography-mass spectrometry/mass spectroscopy performed at a commercial laboratory (Attoquant Diagnostics, Vienna, Austria), as previously described (4,25). Briefly, samples were spiked with a stable isotope labelled internal standard for each angiotensin after *ex vivo* equilibration and analytes were extracted using C18-based solid- phase extraction. Extracts samples were analyzed using mass spectrometry analysis using a reversed- analytical column (Acquity UPLC C18, Waters) operating in line with a XEVO TQ-S triple quadrupole mass spectrometer (Waters Xevo TQ/S, Milford, MA) in multiple reaction monitoring mode. Internal standards were used to correct for analyte recovery across the sample preparation procedure in each individual sample. Analyte concentrations were calculated from integrated chromatograms considering the corresponding response factors determined in appropriate calibration curves in serum matrix, when integrated signals exceeded a signal-to-noise ratio of 10. The lower limits of quantification for the analytes in canine serum were 3 pmol/L (Ang I), 2 pmol/L (Ang II), 2,5 pmol/L (Ang III), 2 pmol/L (Ang IV), 2,5 pmol/L (Ang 1-7), and 2 pmol/L (Ang 1-5), respectively (26,27).

### Data Preparation

Pharmacokinetic and pharmacodynamic data were imported into R 4.0.2 (28) for model preparation. To maintain consistency across administration units and measurement units, all concentration data were converted to micromoles per liter. Benazepril HCl and benazeprilat molecular weights were obtained from PubChem (29,30).

### Placebo

Historical control data from a previous study was used in lieu of a placebo group (23). After referencing parameters to this baseline during fit, all placebo simulation response generated was equivalent to the baseline chronobiology function with parameters estimated from the simultaneous modeling of all biomarkers response.

### NLME Modeling

The recorded data (*y_ij_*) were imported into Monolix 20120 R1 (Lixoft, France) and used to estimate population parameters (***μ***) and variance via the stochastic approximation expectation maximization algorithm (SAEM) (31). Individual parameters (◻_*i*_) were determined via the modes of the individual posterior distributions – which were estimated using a Markov-Chain Monte-Carlo (MCMC) procedure. NLME models were written as previously described (**Eq. 1**) (32,33).

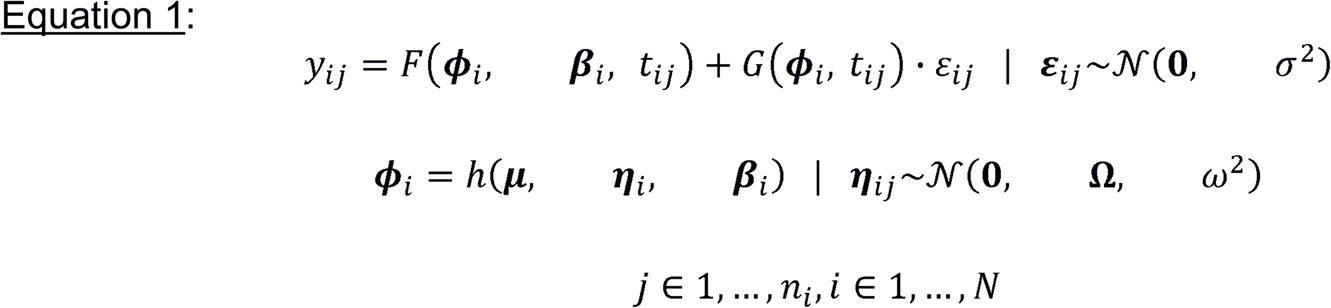

Model predictions (*F(ϕ_i_, β_i_, t_ij_*)) for the *i*^th^ individual at the *j*^th^ timepoint – time (*t_ij_*) – were parameterized using individual parameters and individual covariates (*β_i_*). The residuals were modeled as *G(ϕ_i_, t_ij_)· ε_ij_*, where *G* is an arithmetic combination of proportional and additive error distributions.

Individual parameters were modeled as a function of the population parameters, interindividual variability (***η**_i_*), and individual covariates via the interindividual variation function *h(**μ, η**_i_, **β**_i_)*. Individual variability was modeled with a normal distribution of mean 0, variance-covariance matrix Ω, and variance *ω^2^*. Typically, *h(**μ, η**_i_, **β**_i_)* is a log-normal link function (**Eq. 2**) or a logit-normal link function (**Eq. 3**) in those cases where *ϕ_i_* is bounded to be between 0 and 1.

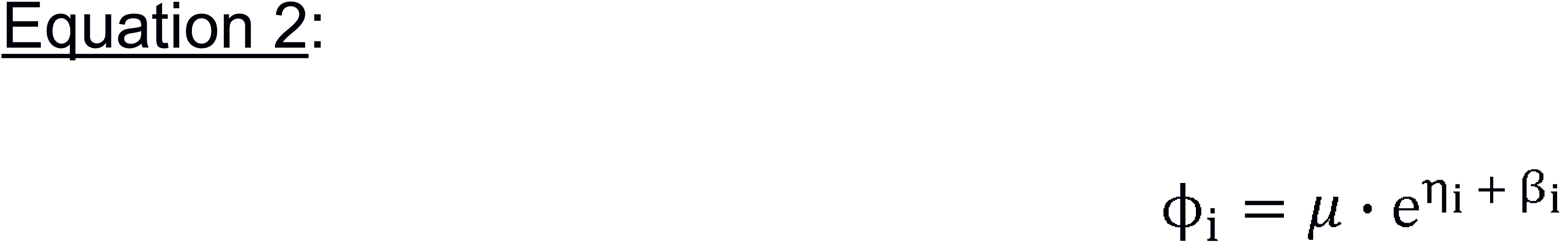

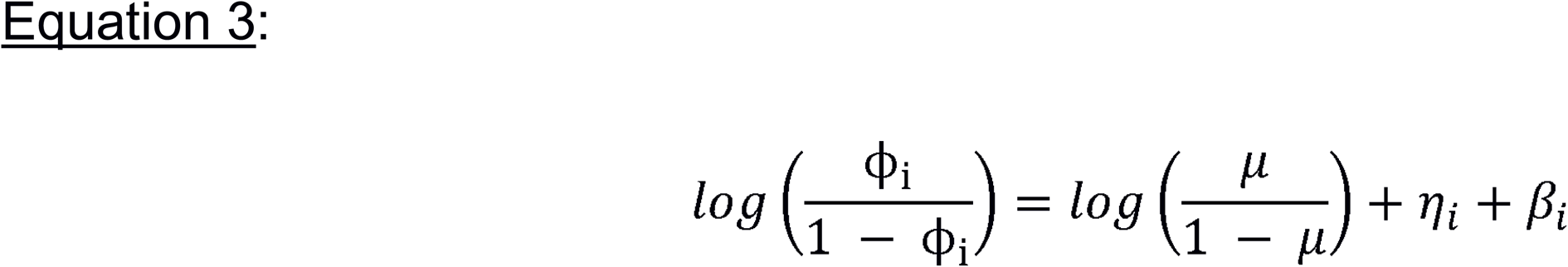

### Model Building

The systems pharmacology model was built in two consecutive phases. First, a largely empirical PKPD model with minimum parameterization was built to capture basic biological variations in the data. In practice, this entailed fitting a basic 1-, 2-, or 3-compartment model to the pharmacokinetics of benazeprilat, and then linking the PK to the RAAS biomarkers concentrations via various indirect and direct response models. Whenever possible we opted for direct over indirect effects models, and fewer compartments, to reduce the number of total estimated parameters during model fit.

Then, in an iterative fashion, model components were replaced with more mechanistic structures. To do this, we modeled the cascade of peptides which define the *alternative* and *classical* RAAS pathways. We also tested whether we could expand the mathematical model to include important biological systems such as the clearance of angiotensins via the liver vs. kidneys, non-specific plasma binding, first-pass metabolism, and site-specific metabolism. Some components and parameters of the model structure were arbitrarily fixed to literature or exploratory values to preserve fidelity to relevant biological systems. For example, our model equations were rewritten so that the production of AngII was always one-to-one proportional with catalysis of AngI via the angiotensin converting enzyme. The final model was refined through various arithmetic simplifications and parameter search optimizations to improve precision of parameter estimates as much as possible without comprising fit to experimental data. The significance of bodyweight, sex, sodium intake, and benazepril dose on parameters estimates was further evaluated using the automated Pearson’s correlation test and ANOVA method as implemented in Monolix 2020 R1.

### Model Evaluation

Experimental data were collated and imported into the 2020R1 Monolix Suite for data exploration, model development and evaluation. Quality of fit was evaluated using standard goodness-of-fit diagnostics (e.g., observed vs. predictions, scatter plot of residuals), as well as numerical summaries of fit, such as the corrected Bayesian information criteria (BICc). Precision of parameter estimates were determined using residual standard error (RSE%).

Normalized prediction distribution errors (NPDEs) are recommended for evaluating model misspecification when study design is heterogenous with respect to dosing groups (34). In this study, we had several different study groups under varying dosing schedules; therefore, NPDEs were chosen to determine quality of fit over more conventional visual predictive checks. In practice, NPDEs evaluate the percentage of the distribution of predictions at each mean prediction under the observation, thus forming a heterogenous uniform distribution. Therefore, an inverse cumulative distribution function is applied to each value to obtain a probability density function vs. population prediction.

### Programming Tools and Simulations

All programming relevant to the simulation application was written in R v4.0.2 (28). Models were translated from Mlxtran (Monolix Suite 2020R1) to the domain specific language, and R package, Odin v1.2.1 (35) to simulate clinical trials. Odin provides an interface for the ODE solvers, and R packages, deSolve v1.30 (36) and dde v1.0.1 (37). Odin was used because of its superior ability to solve large systems of ODEs over other R packages.

Finally, the application for simulating clinical trials was built in Shiny v1.6.0. Shiny is an R package that automatically generates HTML applications from R code. All Shiny applications are designed to (1) generate an HTML-based graphical user interface (GUI) that allows users to interact with the computer hosting the Shiny application, called the *server*, and (2) execute R code on the server based on the user’s interactions with the GUI. To facilitate the use of our simulator, a user-friendly GUI was developed that allows to specify modalities for a clinical trial simulation in R (i.e., defining parameters such as dosage, dosing interval, size of trial) on a website server.

## Acknowledgements

Agnes Bourgois-Mochel (1) and Oliver Domenig (2)

(1) Veterinary Clinical Sciences, Iowa State University, 50011-1250 Ames, IA (USA); (2) Attoquant Diagnostics, Vienna (Austria)

