## Supplementary material for "*Breakthrough*: A First-In-Class Virtual Simulator for Dose Optimization of ACE Inhibitors in Veterinary Cardiology": S1 Table

Ang(1-7)_fr_	≝	amount of angiotensin (1-7) in free circulation compartment

Ang(1-7)_ts_	≝	amount of angiotensin (1-7) in tissue compartment

AngIII_pl_	≝	amount of angiotensin III in plasma compartment

AngIII_rn_	≝	amount of angiotensin III in renal compartment

AngIII_ts_	≝	amount of angiotensin III in tissue compartment

AngIV_pl_	≝	amount of angiotensin IV in plasma compartment

AngIV_rn_	≝	amount of angiotensin IV in renal compartment

AngIV_ts_	≝	amount of angiotensin IV in tissue compartment

δ_24hr_	≝	scale of renin production variance due to chronobiology

ε_Ang(1-7)_	≝	proportional measurement error of angiotensin (1-7)

ε_AngI_	≝	proportional measurement error of angiotensin I (1-10)

ε_AngII_	≝	proportional measurement error of angiotensin II (1-8)

ε_AngIII_	≝	proportional measurement error of angiotensin III (2-8)

ε_AngIV_	≝	proportional measurement error of angiotensin IV (3-8)

ε_benazeprilat_	≝	proportional measurement error of benazeprilat

E_fr_	≝	amount of ACE in free circulation compartment

E_ts_	≝	amount of ACE-bound benazeprilat in tissue compartment

EI_fr_	≝	amount of ACE-bound benazeprilat in free circulation compartment

EI_ts_	≝	amount of ACE-bound benazeprilat in tissue compartment

ES_fr_	≝	amount of ACE-bound angiotensin I (1-10) in free circulation compartment

ES_ts_	≝	amount of ACE-bound angiotensin I (1-10) in free tissue compartment

F_bio_	≝	effective bioavailability of benazepril

f_CT_(t)	≝	effect of chronobiology on the production rate of the angiotensin I (1-10)

I_1abs, Ifr_	≝	fraction of benazepril absorbed from depot to free circulation compartment

I_1abs, Ipr_	≝	fraction of benazepril absorbed from depot to pre-circulation compartment

I_fr_	≝	amount of benazeprilat in free circulation compartment

I_ns_	≝	amount of benazeprilat bound via non-specific interactions in blood compartment

I_pr_	≝	amount of benazeprilat in pre-circulation compartment

I_ts_	≝	amount of benazeprilat in tissue compartment

k_–3_	≝	rate of ACE-benazeprilat association *in vivo*

k_–1_	≝	rate of ACE-angiotensin I (1-10) association *in vivo*

k_1_	≝	rate of ACE-angiotensin I (1-10) disassociation *in vivo*

k_2_	≝	rate of production of angiotensin II (1-8) from ACE-angiotensin I (1-10) *in vivo*

k_3_	≝	rate of ACE-benazeprilat disassociation *in vivo*

ka	≝	zero-order analog absorption rate of benazepril

ka_1_	≝	first-order analog absorption rate of benazepril

k_Cl, Ang(1-7)_	≝	rate of clearance of angiotensin (1-7) from free circulation compartment; defined as

Cl_Ang(1-7)_/V_fr_

k_Cl, I_	≝	rate of clearance of benazeprilat from free circulation compartment; defined as Cl_I_/V_fr_

k_Cl, P_	≝	rate of clearance of angiotensin II (1-8) from free circulation compartment; defined

as Cl_P/_V_fr_

k_Cl, S_	≝	rate of clearance of angiotensin I (1-10) from free circulation compartment; defined

as Cl_S_/V_fr_

k_Cl, AngIII_	≝	rate of clearance of angiotensin III (1-8) from free circulation compartment; defined

as Cl_AngIII_/V_fr_

k_Cl, AngIV_	≝	rate of clearance of angiotensin I (1-10) from free circulation compartment; defined

as Cl_AngIV_/V_fr_

k_fr, ns_	≝	rate of transfer from free circulation compartment to non-specifically bound

interaction compartment; defined as Q_fr, ns_/V_fr_

k_fr, ts_	≝	rate of transfer from free circulation compartment to tissue compartment defined as

Q_fr, ts_/V_fr_

k_I, 1-7_	≝	overall in vivo conversion rate of angiotensin I (1-10) to angiotensin (1-7)

k_II, 1-7_	≝	overall in vivo conversion rate of angiotensin II (1-8) to angiotensin (1-7)

k_II, III_	≝	overall in vivo conversion rate of angiotensin II (1-8) to angiotensin III (2-8)

k_III, IV_	≝	overall in vivo conversion rate of angiotensin III (2-8) to angiotensin IV (3-8)

k_ns, fr_	≝	rate of transfer from non-specifically bound interaction compartment to free

circulation compartment; defined as Q_fr, ns_/V_ns_

k_pl, rn_	≝	rate of transfer from plasma compartment to renal compartment, both within free

circulation compartment; defined as Q_rn_/V_pl_

k_pl, ts_	≝	rate of transfer from plasma compartment within free circulation compartment to

tissue compartment; defined as k_fr, ts ∙_ (V_pl_/V_fr_)

k_rn, pl_	≝	rate of transfer from renal compartment to plasma compartment, both within free

circulation compartment; defined as Q_rn_/V_rn_

k_rn, ts_	≝	rate of transfer from renal compartment within free circulation compartment to

tissue compartment; defined as k_fr, ts ∙_ (V_rn_/V_fr_)

k_ts, fr_	≝	rate of transfer from tissue compartment to free circulation compartment; defined as

Q_fr, ts_/V_ts_

k_ts, pl_	≝	rate of transfer from tissue compartment to plasma compartment within free

circulation compartment; defined as k_ts, fr ∙_ (V_pl_/V_fr_)

k_ts, rn_	≝	rate of transfer from tissue compartment to renal compartment within free

circulation compartment; defined as k_ts, fr ∙_ (V_rn_/V_fr_)

P_fr_	≝	amount of angiotensin II (1-8) in free circulation compartment

PRA	≝

P_ts_	≝	amount of

r_S_	≝

S_fr_	≝	amount of angiotensin I (1-10) in free circulation compartment

S_ts_	≝	amount of angiotensin I (1-10) in tissue compartment

V_fr_	≝	volume of distribution associated with free circulation compartment; defined as V_pl_ +

V_rn_

V_pl_	≝	volume of distribution associated with plasma compartment within free circulation

compartment

V_rn_	≝	volume of distribution associated with renal compartment within free circulation

compartment

V_ts_	≝	volume of distribution associated with tissue compartment

Y_Ang(1-7)_	≝	predicted concentration of angiotensin (1-7) in free circulation compartment

Y_AngI_	≝	predicted concentration of angiotensin I (1-10) in free circulation compartment

Y_AngII_	≝	predicted concentration of angiotensin II (1-8) in free circulation compartment

Y_AngIII_	≝	predicted concentration of angiotensin III in free circulation compartment

Y_AngIV_	≝	predicted concentration of angiotensin IV in free circulation compartment

Y_benazeprilat_	≝	predicted concentration of benazeprilat in free circulation compartment
